# An efficient artificial esterase with a dynamic conformation via conformational engineering

**DOI:** 10.1101/2025.06.26.661850

**Authors:** Yan Wang, Yi Cao, Yuan-Yuan Liu, Yuye Cao, Tiange Gao, Jiewen Deng, Tongtong Zhou, Haifang Wang, Aoneng Cao

## Abstract

The catalytic turnover number (*k*_*cat*_) of current enzyme mimics and de novo designed enzymes is still orders of magnitude lower than that of natural enzymes, because mimicking the fast-dynamic feature of enzyme active centers remains a great challenge. Herein, we created a gold nanoparticle-based artificial esterase, Goldenzyme, by conformational engineering to reconstruct dynamic active centers resembling that of α-chymotrypsin. NMR characterization shows that Goldenzyme possesses a fast-dynamic feature, about 6-fold faster than typical α-helix. Therefore, Goldenzyme efficiently hydrolyzes *p*-nitrophenyl acetate with a net *k*_*cat*_ of 6.37 s^−1^ per active center, more than 5-fold of that of α-chymotrypsin (1.19 s^−1^), and Goldenzyme can even hydrolyze some non-activated esters that α-chymotrypsin cannot. Our conformational engineering approach demonstrates the possibility of creating artificial enzymes that surpass natural ones.

## Main Text

Enzymes are miraculous catalysts essential for life. What makes enzymes so marvelous is that they can efficiently catalyze difficult reactions by precisely arranging multiple residues in 3D space and making these residues act synergistically in a concerted manner, as demonstrated by hydrolases, the largest class of enzymes. Typical hydrolases, such as esterases, contain a binding site for the substrate, a catalytic triad (or dyad), and an oxyanion hole, forming an efficient and dynamic charge-relay network to hydrolyze substrates (*1, 2*). Although the catalytic triad (or dyad) and oxyanion hole have been extensively mimicked by different approaches from molecules and molecular assembles (*3–12*), peptide aggregates (*13–16*), dendrimers (*17, 18*), surface-functionalized nanoparticles (NPs) (*19–22*), imprinted polymers (*23–25*), to de novo designed enzymes (*26–31*), and despite the fact that nowadays’ AI-based computational design can produce de novo hydrolases with precisely-defined active centers as supported by x-ray structures (*31*), there is still a huge gap in catalytic activity between current hydrolase mimics and natural hydrolases. Without metal ion cofactors, current artificial hydrolases and de novo designed hydrolases typically show catalytic turnover numbers (*k*_*cat*_) several orders of magnitude lower than those of natural hydrolases even for the hydrolysis of easily hydrolysable activated esters, and show almost no or a very limited catalytic activity for the hydrolysis of non-activated esters (*31– 34*).

Apparently, there might be certain important features of enzymes being missing in current enzyme-mimicking and design. One important feature of enzymes is their fast-dynamic conformational change during catalytic processes, basically, “*enzymes at work are enzymes in motion*” (*35–37*). However, this fast-dynamic feature has not yet been considered in enzyme-mimicking and design, and it is evident that, even for the current powerful AI, mimicking this feature remains a grand challenge (*38,39*). Herein, we report a special conformational engineering (CE) approach to create a gold NP (AuNP)-based efficient artificial esterase, denoted as Goldenzyme, through reconstruction dynamic active centers resembling that of α-chymotrypsin on AuNPs. The fast-dynamic feature endows Goldenzyme a highly efficient catalytic activity for the hydrolysis of esters with a *k*_*cat*_ higher than that of α-CT.

## Results

### Rationale for the design of Goldenzyme

Our goal is to construct a catalytic triad, an oxyanion hole, and a substrate binding site on AuNP from scratch, by a special CE approach to precisely arrange the key catalytic amino-acid residues corresponding to those of α-CT with a dynamic feature. Noticing that the key catalytic residues in many hydrolases are arranged roughly in a line (Fig. 1A–C), we reasoned that we could incorporate these key residues in a single short peptide, and arrange them roughly in a line on the same side of an α-helical conformation (Fig. 1D,E).

**Fig. 1.**
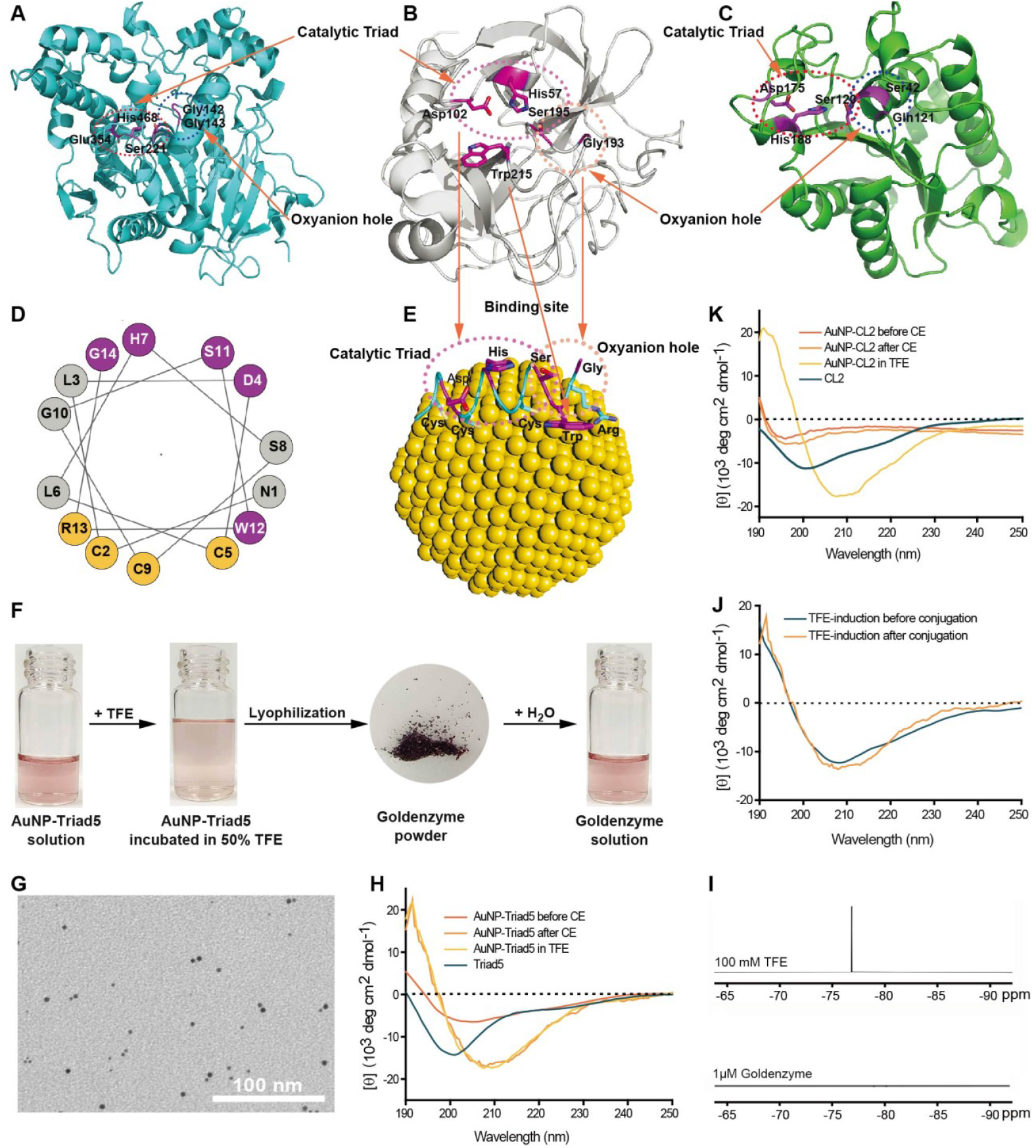
Design, synthesis, and characterization of Goldenzyme. (**A**) Catalytic triads and oxyanion hole in human liver esterase (PDB id: 1mx5). (**B**) Key catalytic residues of α-CT (PDB id:1cbw): Asp102, His57, and Ser195 form the catalytic triad; Ser195 and Gly193 form the oxyanion hole; and the binding site for substrates is located around Trp215. (**C**) Catalytic triad and oxyanion hole in Fusarium solani cutinase (PDB id: 1cex). (**D**) Sequence of Triad5 arranged in an α-helical conformation. The catalytic key residues are highlighted in magenta, the residues highlighted in orange are intended to anchor to the surface of AuNPs. (**E**) Model of Goldenzyme. For clarity, only one Triad5 peptide is shown. (**F**) Process of the post-synthesis CE used to produce Goldenzyme. (**G**) TEM image of Goldenzyme. (**H**) CD spectra of free Triad5 peptide, the AuNP-Triad5 before CE, the AuNP-Triad5 in 50% TFE, and the AuNP-Triad5 after CE (TFE had been removed by lyophilization). (**I**) ^19^F NMR spectra of 100 mM TFE and 1 μM Goldenzyme. (**J**) Far-UV CD spectra of AuNP-Triad5 with TFE induction before or after Triad5 conjugation. (**K**) Far-UV CD spectra of free CL2 peptide, the AuNP-CL2 before CE, the AuNP-CL2 in 50% TFE, and the AuNP-CL2 after CE (TFE had been removed by lyophilization).

Moreover, noticing that many key residues of catalytic triads and oxyanion holes in hydrolases locate in flexible loop fragments (Fig. 1A–C), we hypothesized that the backbone movement of these fragments is essential for the efficient catalytic activity of hydrolases. Therefore, we plan to keep every coil of the helix flexible, i.e., in an α-helix-like but unstable conformation, so that the movement of the coils can mimic that of the corresponding fragments of hydrolases. To simultaneously meet these two contradictory requirements (helical and flexible), we design a peptide, Triad5 (Fig. 1D and table S1). Triad5 has a certain degree of preference for

α-helix (containing two Leu residues (L3 & L6) and an N-capping residue (N1) (*40*)), but cannot form stable helical conformation by itself. When adopting an α-helix-like conformation, an Asp (D4), a His (H7), and a Ser (S11) will locate on the same side but different coils of the helix, forming the catalytic triad; and three Cys (C2, C5, C9) and an Arg (R13) located on the opposite side of the helix will help to anchor each of the three coils onto the AuNP via Au-S bonds and the strong interaction of the guanidino group of Arg with the AuNP (Fig. 1D,E). Restricted by its neighbor Arg (R13), the C-terminal Gly (G14) forms the oxyanion hole together with the Ser (S11). And the Trp (W12) residue is the designed binding site for substrates. Thus, all the key catalytic residues of α-CT are incorporated in the designed Triad5 peptide (Fig. 1B,E).

The key step for the synthesis of Goldenzyme is to finely tune Triad5 on AuNPs into an approximately helical yet flexible conformation. This tricky problem is solved by a special CE technique, which turns out to be surprisingly simple. Basically, we first synthesize the AuNP-Triad5 conjugate, then induce Triad5 on the AuNP into an α-helical conformation by adding trifluoroethanol (TFE), a widely-used α-helical conformation inducer (*41*), and finally remove TFE by lyophilization (Fig. 1F). As we demonstrated previously (*42–48*), the distance between Au-S bonds, i.e. the anchoring positions of Cys residues on AuNPs, will be adjusted accordingly to the induced conformation of peptides, due to the mobility of the surface atoms of AuNPs (*42,49– 52*) and the conformation-preference of the conjugated peptides (*53,54*). However, after removing TFE, the conformation of Triad5 on the AuNP would not remain in stable α-helix, because the weak preference of the peptide for the α-helical conformation. Yet, due to the strong Au-S bonds as well the interaction between Arg and the surface of AuNPs, the anchoring positions of the Cys and Arg residues on AuNPs would be largely remained, thereby keeping the key residues flexible in the right proximity for dynamic and cooperative catalysis.

### Conjugation of Triad5 onto AuNPs and the CE technique

Triad5 was conjugated onto citrate-capped AuNPs (average diameter is about 3.6 nm, Fig. 1G and fig. S1). As we previously demonstrated (*42–48*), the peptide density (the average number of conjugated peptides per NP) on the surface of AuNPs plays an important role in maintaining the right conformation of the conjugated peptides, therefore Triad5 was conjugated onto AuNPs with different densities to find the optimal one. Since the conjugation is very efficient (fig. S2), the pre-determined peptide density could be achieved by mixing quantified AuNPs and peptides.

To induce Triad5 of the AuNP-Triad5 conjugate into α-helical conformation, TFE was added to a final concentration of 50% (v/v), and then the sample was incubated at different temperatures for different periods of time. Afterwards, TFE was removed from the sample by lyophilization, resulting in a dark purplish-red power, which can be easily re-dispersed in water (Fig. 1F).

Circular dichroism (CD) spectra show that free Triad5 is basically in random coil conformation, and Triad5 of the as-conjugated AuNP-Triad5 (i.e., the AuNP-Triad5 before CE) is also largely unstructured or partially structured (Fig. 1H). After incubation in 50% TFE, Triad5 of the AuNP-Triad5 is induced into α-helical conformation. Remarkably, the CD spectrum of the AuNP-Triad5 after CE (TFE had been removed by lyophilization, and the residual TFE was estimated to be less than 1 mM (Fig. 1I), which has no α-helix inducing effect) is almost the same as that of the AuNP-Triad5 in TFE (Fig. 1H). Interestingly, if Triad5 is pre-incubated in 80% TFE and then conjugated to AuNP in 50% TFE to ensure that Triad5 is in an α-helical conformation, after lyophilization and re-dispersion in aqueous solutions, the resulted AuNP-Triad5 (TFE-induction before conjugation) has the same CD spectrum as the above AuNP-Triad5 after CE (TFE-induction after conjugation) (Fig. 1J). All these results prove that the CE treatment can indeed tune the conformation of the conjugated Triad5 into an α-helical conformation.

We also conjugated AuNPs with another peptide, CL2, which is adopted from the CDR3 fragment of an anti-lysozyme antibody (*45,55*) and lacks the catalytic His residue and the oxyanion hole. CL2 is in loop conformation in the original antibody and on the surface of AuNPs (*45,55*). Therefore, the solution of AuNP-CL2 is a perfect blank for AuNP-Triad5. As shown in Fig. 1K, CL2 of AuNP-CL2 could also be induced into α-helical conformation in 50% TFE, however, it reverted to the loop conformation after removing TFE, highlighting the importance of the conformational preference of the designed peptide (*53,54*).

The catalytic activity of AuNP-Triad5 and the CE effect were evaluated by measuring the catalytic hydrolysis of *p*-nitrophenyl acetate (*p*-NPA), a common substrate for the assays of hydrolases and hydrolase-mimics due to its easy quantification of the catalytic mechanism and kinetics (*56*). Since the AuNP-Triad5 after CE shows a very high efficiency for the hydrolysis of *p*-NPA, we had to use a stopped-flow instrument to carry out and monitor the fast reaction (fig. S3). According to the normalized peptide-based initial catalytic rates, the optimal CE condition was determined as 50% (v/v) TFE for 24 hours at 45 °C (Fig. 2A and fig. S4). The slightly decreased activity after prolonged incubation in TFE at 45 °C is because TFE could induce amyloid aggregation of proteins and peptides, especially at elevated temperatures (*41*). Similarly, the optimum peptide density was determined to be 20 Triad5 per AuNP (Fig. 2B, fig. S5, and table S2). Interestingly, TFE-induction, whether before or after conjugation, results in the same catalytic activity (Fig. 2C). This result further confirme the CE effect. So, hereafter, the AuNP-20Triad5 after CE (simplified as AuNP-Triad5 after CE) is referred to as Goldenzyme.

**Fig. 2.**
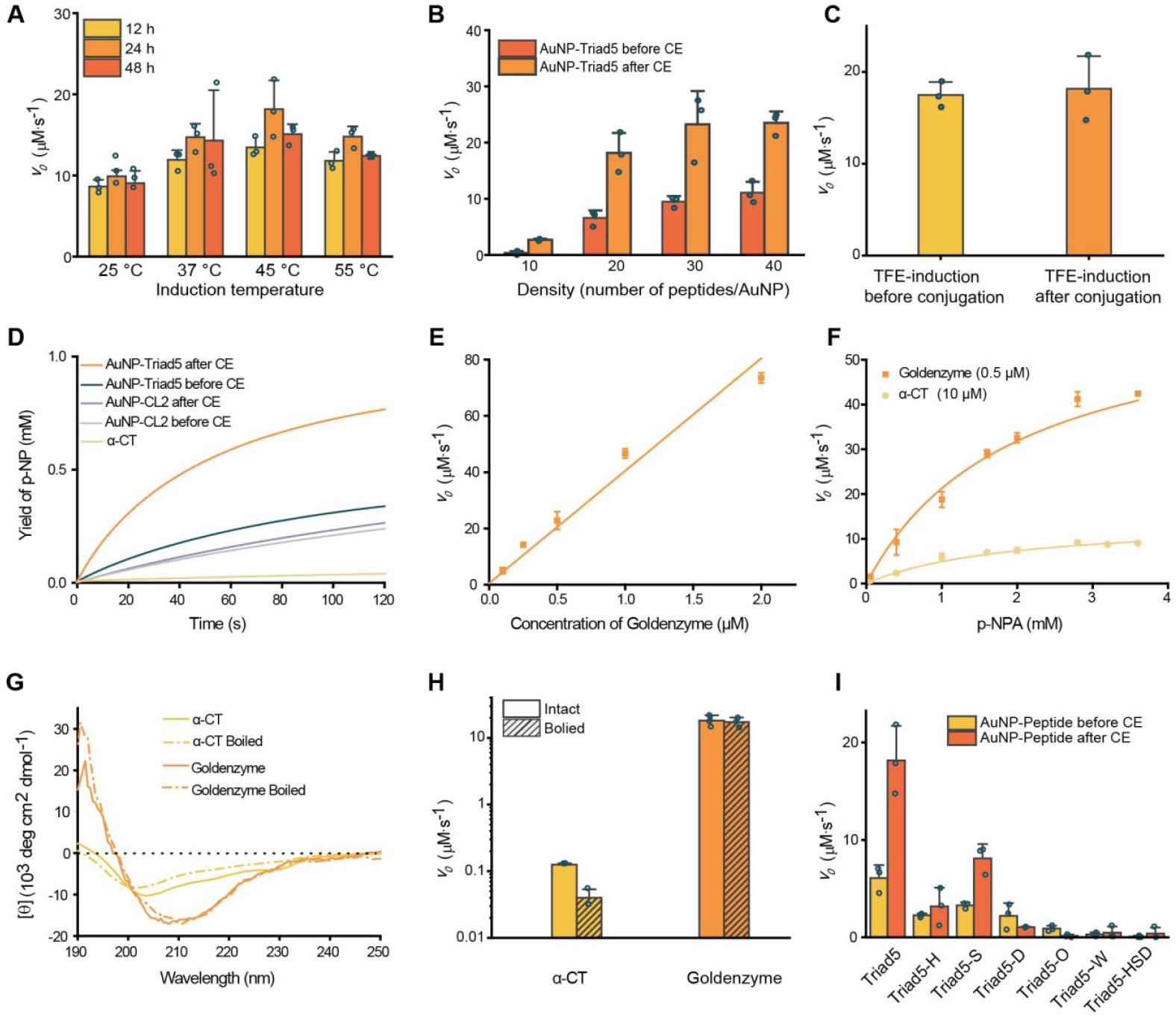
Catalytic hydrolysis of *p*-NPA. (**A**) Effect of CE incubation temperature and time on the initial rate of catalytic hydrolysis of *p*-NPA by the AuNP-Triad5 after CE. (**B**) Effect of peptide density on the initial rate of catalytic hydrolysis of *p*-NPA by the AuNP-Triad5 after CE. (**C**) Catalytic activity of Goldenzyme prepared by the TFE induction before or after conjugation. (**D**) Hydrolysis of *p*-NPA catalyzed by different catalysts. (**E**) Correlation between the concentration of Goldenzyme and its catalytic activity. (**F**) Initial hydrolysis rates of *p*-NPA at different concentrations catalyzed by 0.5 μM Goldenzyme (orange squares) and 10 μM α-CT (yellow circles) in the same buffer. Data were fitted to the Michaelis–Menten equation. (**G**) Far-UV CD spectra of α-CT (yellow) and Goldenzyme (orange) before (solid lines) and after (dashed lines) being boiled at 100 °C for 1 hour. (**H**) Initial hydrolysis rates of *p*-NPA catalyzed by enzymes before and after being boiled at 100 °C for 1 hour. (**I**) Initial hydrolysis rates of *p*-NPA catalyzed by the AuNP-Triad5 and its mutants before and after CE. Conditions for the hydrolysis: [Goldenzyme/mutants/α-CT] = 0.5 μM (except that in (E, F)), [*p*-NPA] = 1 mM (except that in (F)), [acetonitrile] = 2% (v/v), [PB] = 40 mM, pH = 7.0, *T* = 25 °C. Error bars indicate SDs of triplicate experiments. The catalytic rates of Goldenzyme and its mutants in (B,I) have subtracted that of the corresponding AuNP-CL2 (before or after CE).

Goldenzyme powder is readily dissolved in aqueous solutions and shows a much higher catalytic activity than the free Triad5 and the AuNP-Triad5 before CE (Fig. 2D and fig. S3), and the catalytic activity of Goldenzyme is concentration-dependent (Fig. 2E and fig. S6), highlighting the importance and success of CE. Remarkably, with ∼20 active sites in a similar size to natural enzymes, Goldenzyme shows a much higher catalytic rate for the hydrolysis of *p*-NPA than α-CT (Fig. 2F), and even comparable to porcine liver esterase (fig. S7).

### Catalytic efficiency of Goldenzyme for the hydrolysis of p-NPA

Although *p*-NPA is the most widely used substrate for hydrolase-mimicking studies, it is known to be very unstable, and many factors can accelerate its hydrolysis. To accurately quantify the net catalytic activity of Goldenzyme, it is necessary to subtract proper blanks.

Besides pH buffer, both the AuNP core and the high concentration of accompanying salts (resulting from the lyophilization) contributes for the observed activity of AuNP-peptide conjugates for the hydrolysis of *p*-NPA, and must therefore be subtracted. As shown in fig. S3A, the AuNPs per se significantly accelerated the hydrolysis of *p*-NPA.

To quantify the contribution of the high salts in AuNP-peptide conjugate solutions, we ultrafiltrated the AuNP-peptide (before CE) solution and used the filtrate as the buffer before CE. And the buffer after CE was prepared by treating the buffer before CE with TFE in the same way as the CE treatment of Goldenzyme, so the buffer after CE contained the same concentration of all solutes including trace TFE as in the Goldenzyme solution. As expected, *p*-NPA in the buffer before CE shows a higher hydrolysis rate than in the common PB buffer (fig. S3 and fig. S8A,B), and the hydrolysis rate in the buffer after CE is higher than in the buffer before CE, indicating that the residual TFE in the buffer still has certain effects.

To accurately quantify the net catalytic activity of Goldenzyme, the AuNP-CL2 after CE is a perfect blank to be subtracted, because it literally contains all the influencing factors for Goldenzyme, including buffer (pH and ionic strength), trace TFE, AuNPs, and the same number of peptides (but non-active). As shown in fig. S3B, the AuNP-CL2 after CE shows a similar (slightly higher) activity compared to the AuNPs after CE, and the AuNP-CL2 before CE also shows a similar activity to the AuNPs before CE. Notably, the difference in activity between

AuNP-CL2 before and after CE equals to that between AuNPs before and after CE, indicating that this difference is simply the difference between the buffers and thus can be reliably subtracted.

After subtraction of the corresponding blank, the initial catalytic rates (*v*_0_) of Goldenzyme and the controls were fitted to the Michaelis–Menten equation (Fig. 2F, Table 1 and fig. S9). And for a fairer comparison, the hydrolysis by α-CT was also carried out in the buffer after CE, and the data were also fitted to the Michaelis–Menten equation (Fig. 2F and Table 1). As can be seen from Table 1, the free Triad5 peptide in the buffer after CE has a tiny *k*_*cat*_ of 0.0057 s^−1^ (fig. S9A). However, after being conjugated to AuNPs, the AuNP-Triad5 before CE shows a substantially increased net *k’*_*cat*_ of 37.3 s^−1^ (Table 1 and fig. S9B). After CE, the net *k’*_*cat*_ of the AuNP-Triad5 after CE (Goldenzyme) further rises to 127.4 s^−1^ (Table 1 and Fig. 2F), even higher than the porcine liver esterase, which has a *k*_*cat*_ of 106 s^−1^ (fig. S7C) (*57*).

**Table 1.** Kinetic parameters for catalyzing the hydrolysis of p-NPA.

| Catalyst | $k_{cat}$<br>($\text{s}^{-1}$)<sup>a</sup> | $k'_{cat}$<br>($\text{s}^{-1}$)<sup>b</sup> | $K_m$<br>(mM) | $k_{cat}/K_m$<br>($\text{M}^{-1}\cdot\text{s}^{-1}$) | $(k_{cat}/K_m)'$<br>($\text{M}^{-1}\cdot\text{s}^{-1}$)<sup>e</sup> |
| --- | --- | --- | --- | --- | --- |
| Free Triad5 | $0.0057 \pm 0.0007$ | / | $1.2 \pm 0.4$ | 4.8 | / |
| AuNP-Triad5 before CE | $53.2 \pm 23$ | $37.3^c$ | $2.2 \pm 1.3$ | $2.4 \times 10^4$ | $1.6 \times 10^4^f$ |
| AuNP-Triad5 after CE (**Goldenzyme**) | $147 \pm 14$ | $127.4^d$ | $2.0 \pm 0.5$ | $7.4 \times 10^4$ | $6.7 \times 10^4^g$ |
| AuNP-Triad5-D after CE | $25.2 \pm 4.1$ | $5.6^d$ | $3.3 \pm 0.6$ | $7.6 \times 10^3$ | $1 \times 10^2^g$ |
| AuNP-Triad5-H after CE | $55.6 \pm 35$ | $36.0^d$ | $3.3 \pm 2.7$ | $1.7 \times 10^4$ | $9.5 \times 10^3^g$ |
| AuNP-Triad5-S after CE | $68.2 \pm 8.1$ | $48.6^d$ | $2.5 \pm 0.6$ | $2.7 \times 10^4$ | $1.9 \times 10^4^g$ |
| AuNP-Triad5-HSD after CE | $21.6 \pm 7.1$ | $2.0^d$ | $3.0 \pm 2.0$ | $7.2 \times 10^3$ | $-3 \times 10^2^g$ |
| AuNP-Triad5-O after CE | $28.8 \pm 30$ | $9.2^d$ | $4.0 \pm 5.5$ | $7.2 \times 10^3$ | $-3 \times 10^2^g$ |
| AuNP-Triad5-W after CE | $17.1 \pm 3.6$ | $-2.5^d$ | $0.83 \pm 0.3$ | $2.1 \times 10^4$ | $1.4 \times 10^4^g$ |
| AuNP-CL2 before CE | $15.9 \pm 4.3$ | / | $2.1 \pm 0.6$ | $7.6 \times 10^3$ | / |
| AuNP-CL2 after CE | $19.6 \pm 3.2$ | / | $2.6 \pm 0.7$ | $7.5 \times 10^3$ | / |
| α-CT<sup>h</sup> | $1.19 \pm 0.68$ | / | $1.3 \pm 0.8$ | $9.1 \times 10^2$ | / |
| α-CT<sup>i</sup> | $0.30 \pm 0.01$ | / | $0.36 \pm 0.01$ | $8.3 \times 10^2$ | / |

<sup>a</sup> when calculating $k_{cat}$ , only the value of the corresponding buffer has been subtracted.
<sup>b</sup> $k'_{cat}$ indicates the net turnover number after subtracting that of AuNP-CL2 (before or after CE).
<sup>c</sup> value after subtracting the $k_{cat}$ of AuNP-CL2 before CE.
<sup>d</sup> value after subtracting the $k_{cat}$ of AuNP-CL2 after CE.
<sup>e</sup> $(k_{cat}/K_m)'$ indicates the catalytic efficiency ( $k_{cat}/K_m$ ) after subtracting that of AuNP-CL2 (before or after CE).
<sup>f</sup> value after subtracting the $k_{cat}/K_m$ of AuNP-CL2 before CE.
<sup>g</sup> value after subtracting the $k_{cat}/K_m$ of AuNP-CL2 after CE.
<sup>h</sup> measured with $10 \mu\text{M}$ α-CT in the same buffer as that of Goldenzyme.
<sup>i</sup> measured with $0.5 \mu\text{M}$ α-CT in PB buffer.

Of course, Goldenzyme has an advantage of 20 active sites on the same size of natural enzymes. But even divided by 20, the net *k’*_*cat*_ per active site of Goldenzyme (6.37 s^−1^) is still more than 5 times that of α-CT (1.19 s^−1^) under the same condition (Table 1). By the way, the measured *k*_*cat*_ of α-CT in PB buffer (0.30 s^−1^) (fig. S8C and Table 1) is much lower, but is comparable to the value reported in the literature (*58*), indicating that the effect of the high salts on enzymes is similar, whether they are natural or artificial ones.

Besides its high catalytic activity, Goldenzyme also shows excellent thermal stability due to its stable core, AuNP. As shown in Fig. 2G, the CD spectrum of Goldenzyme does not change after being boiled at 100 °C for 1 hour. As a comparison, the CD spectrum of α-CT undergoes substantial changes after the same thermal treatment. Consequently, α-CT loses most of its catalytic activity after the thermal treatment, while there is almost no decrease in the catalytic activity of Goldenzyme (Fig. 2H and fig. S10).

### Roles of individual key residues of Goldenzyme and the power of CE

To verify the roles of individual key residues in the catalytic efficiency of Goldenzyme, a series of mutants of Triad5 (table S1) were conjugated to AuNPs to produce mutants of Goldenzyme.

As the catalytic activity of these Goldenzyme mutants was significantly reduced (fig. S11), after subtraction of the data of the AuNP-CL2 after CE, the “net” *v*_*0*_ of the mutants could not be fitted to the Michaelis–Menten equation, indicating that each of those key residues is indispensable. To obtain a semiquantitative evaluation, the net catalytic parameters of these mutants were obtained by fitting their original *v*_*0*_ to the Michaelis–Menten equation (Table 1 and fig. S9C–H) to get rough parameters and then subtracting those of the AuNP-CL2 after CE (Table 1 and fig. S9J).

Before the semiquantitative evaluation of the mutation effects of Goldenzyme, it should be noted that even though the catalytic triad is the most important part of natural hydrolases, a single-residue mutation cannot eliminate the catalytic activity of the enzyme completely. In fact, it had been shown that even after all the three residues of the triad of trypsin being mutated simultaneously, the mutant (H57A/D102N/S195A) still showed a catalytic activity (*k*_*cat*_) of approximate 100000 times higher than the hydrolysis without catalysis (*k*_*uncat*_) (*59*). The *k*_*cat*_/*k*_*uncat*_ ratio of this mutant lacking the catalytic triad is even larger than those of many enzyme-mimics.

In addition, although it is normal to see several orders of magnitude change in activity for single mutation of the key residues of natural enzymes, the mutation effect on activity varies greatly depending on hydrolysis conditions. For example, after Asp102 of trypsin was mutated to Asn, the *k*_*cat*_ of the mutant is about 1/10000 of that of the wild type at neural pH, but rises to 6% of that of the wild type at pH 10.2 (*60*).

In the current study, the high concentration of salts has similar effects on the hydrolysis as the increasing pH for the trypsin mutant, especially when the activated ester is used as the substrate. Therefore, the net *k’*_*cat*_ values of single mutants for each of the catalytic triad residues are several times, but not several orders of magnitude, less than that of Goldenzyme (Table 1). Among them, the Triad5-D mutant, whose Asp was mutated to Ala, shows the smallest net *k’*_*cat*_ of 5.6 s^−1^, more than 22 times less than that of Goldenzyme. And the overall catalytic efficiency (*k*_*cat*_/*K*_*m*_) of the Triad5-D mutant is only slightly larger than that of the control (AuNP-CL2 after CE), indicating that Asp is an indispensable catalytic residue. The other two single triad-residue mutants of Goldenzyme also show significantly decreased activity. The net *k’*_*cat*_ of the Triad5-H mutant (His mutated to Ala) is 36.0 s^−1^, and that of Triad5-S mutant (Ser mutated to Ala) is 48.6 s^−1^. A possible explanation for the least decreased *k’*_*cat*_ of Triad5-S mutant is that the remaining Asp and His can still form a catalytic dyad, as that in some enzymes (*61*). This possibility is supported by the fact that the Triad5-S mutant is the only mutant that still shows the CE effect, as its activity significantly increased after CE (Fig. 2I, fig. S11 and table S3), which means that CE is required to correctly fold the Triad5-S peptide on AuNPs and bring its Asp and His to the right positions to form the catalytic dyad.

To further demonstrate the indispensable role of the designed catalytic triad, a mutant with all the three residues being mutated to Ala (table S1) was synthesized. As expected, this AuNP-Triad5-HSD mutant showed a further reduced net *k’*_*cat*_ of 2.0 s^−1^, much less than all three single-residue mutants (Table 1 and fig. S11). Remarkably, this AuNP-Triad5-HSD mutant still showed a slightly higher activity than that of the AuNP-CL2 (a small positive value in Fig. 2I), just like the natural enzyme (*59*), because “the catalytic triad is not the sole source of protease catalytic power, …remaining activity reflects the contribution of the oxyanion hole…” (*2*).

To demonstrate the indispensable role of the designed oxyanion hole, the two C-terminal residues of Triad5 for the oxyanion hole were deleted. As shown in Table 1, this Triad5-O mutant (lacking the oxyanion hole) shows a dramatically decreased net *k’*_*cat*_ of 9.2 s^−1^, more than one order of magnitude lower than that of Goldenzyme (Table 1 and fig. S9G), and its overall catalytic efficiency (*k*_*cat*_/*K*_*m*_) is only comparable to that of the non-active control (AuNP-CL2 after CE) (within experimental error), highlighting the importance of the oxyanion hole for the catalysis.

Last, we investigated the coordination between the catalytic residues and the binding site using the Triad5∼W mutant, where the binding residue Trp (W12) was not deleted, but its position was changed with the N-terminal Asn (N1) (table S1). Without change in the composition of the sequence and the catalytic residues, this residue-swapped mutation results in a very low activity that is comparable to that of the non-active control (AuNP-CL2 after CE) (within experimental error) (Table 1 and fig. S9H). Even though the altered site of Trp results in a higher affinity for the substrate, with a *K*_*m*_ of 0.83 mM, its net overall catalytic efficiency is still much lower than Goldenzyme, indicating that the originally designed binding site mimicking that of α-CT is indeed much better at orientating substrates toward the catalytic center. The observed mutation effect of the Triad5∼W mutant is also very similar to the mutation effect of natural enzymes, where a single mutation at the binding site would dramatically reduce the catalytic activity (*62*).

Overall, all six mutants of Goldenzyme, each representing a key catalytic residue or combination, show remarkable mutation effects that are consistent with our design and the mechanism of natural hydrolases. And most notably, the sum of the activities of AuNP-Triad5-O (lacking the oxyanion hole), AuNP-Triad5∼W (with a mislocated binding site) and AuNP-Triad5-HSD (lacking the triad) is much less than the activity of Goldenzyme (Fig. 2I), clearly demonstrating the synergistic effects of the three catalytic units of Goldenzyme, i.e. the catalytic triad, the binding site and the oxyanion hole, in the same way as natural enzymes.

### Catalytic efficiency of Goldenzyme for the hydrolysis of non-activated esters

As *p*-NPA is an activated ester whose stability is sensitive to solutions (resulting in “sensitive” turnover number without catalysis, *k*_*uncat*_), many claims of the efficiency of esterase-mimics based on the hydrolysis of *p*-NPA (usually with a very large *k*_*cat*_/*k*_*uncat*_ but a very small *k*_*cat*_) could not be reproduced when applied to the hydrolysis of non-activated esters. This phenomenon is even called “*p*-nitrophenyl ester syndrome” (*56*). Therefore, in recent years, hydrolase-mimicking has started to tackle the more challenging target of non-activated esters (*63–66*).

To verify the catalytic efficiency of Goldenzyme for non-activated esters, diethylene glycol terephthalate (DTP) was chosen as the first substrate to investigate both the catalyzed hydrolysis and transesterification at pH 8 (Fig. 3A). As shown in Fig. 3B and figs. S12, S13, Goldenzyme shows a higher activity than α-CT for the hydrolysis of DTP, too. As comparisons, the AuNP-Triad5 before CE, the AuNP-Triad5-HSD after CE, and the AuNP-CL2 after CE show significantly lower activities than Goldenzyme and α-CT (Fig. 3B and figs. S14–S16), clearly demonstrating the important role of the designed catalytic triad and the effect of CE. In addition, Goldenzyme also shows good recyclability for future applications (fig. S17).

**Fig. 3.**
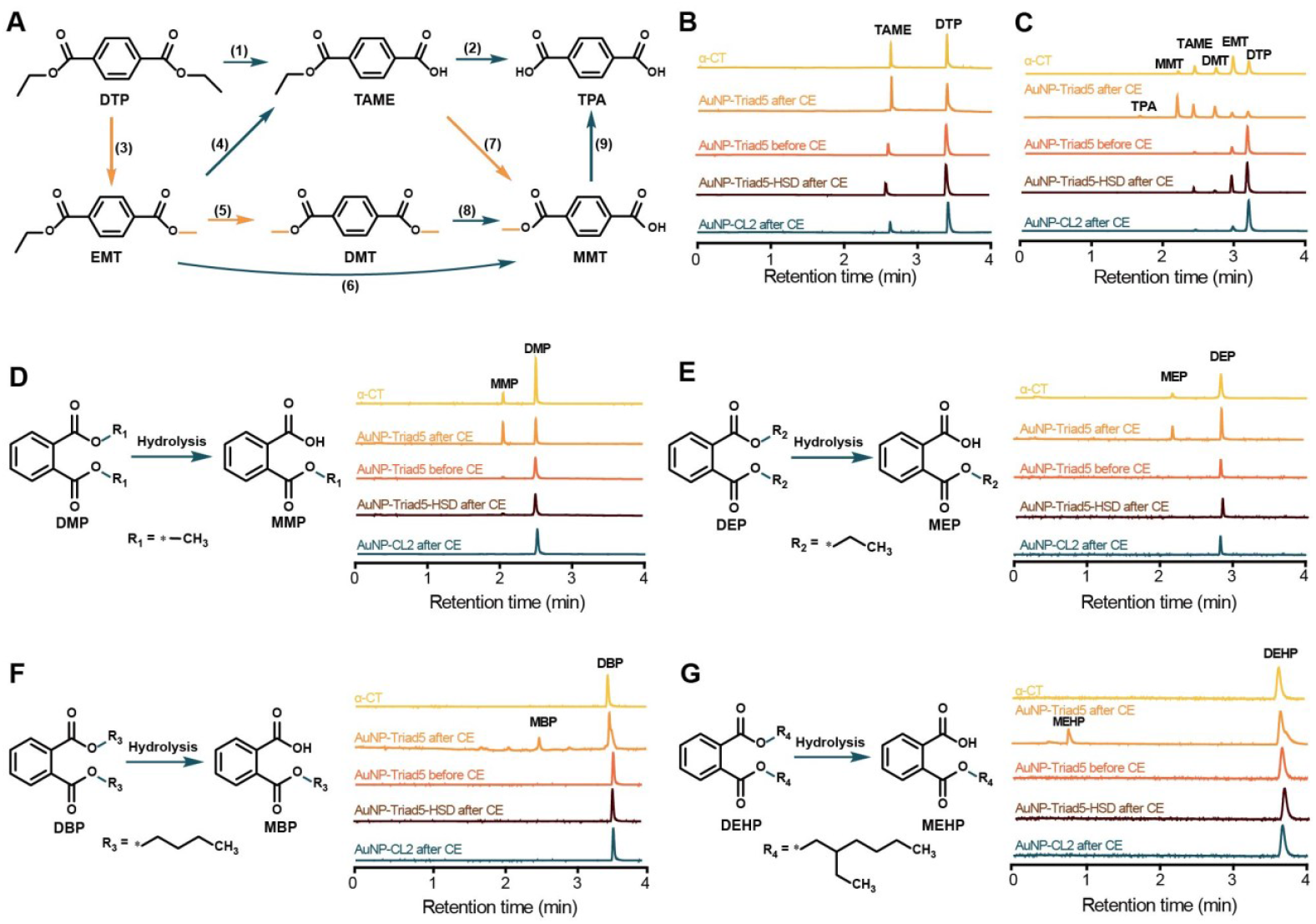
Catalytic hydrolysis of non-activated esters. (**A**) Hydrolysis processes of DTP in the absence/presence of methanol. Blue arrows indicate hydrolysis, orange arrows indicate transesterification. In the absence of methanol, only the process (1) & (2) happen, while all processes happen in the methanol + water solution. (**B**) Species after 48 hours hydrolysis of DTP at 37 °C in the absence of methanol and catalyzed by 0.8 μM α-CT (yellow), the AuNP-Triad5 after CE (Goldenzyme, orange), the AuNP-Triad5 before CE (tangerine), the AuNP-Triad5-HSD after CE (brown) or the AuNP-CL2 after CE (blue). (**C**) Species after 24 hours hydrolysis of DTP at 37 °C in 10% (v/v) methanol solution and catalyzed by 0.8 μM α-CT (yellow), the AuNP-Triad5 after CE (orange), the AuNP-Triad5 before CE (tangerine), the AuNP-Triad5-HSD after CE (brown) or the AuNP-CL2 after CE (blue). (**D**) Species after 48 hours hydrolysis of DMP at 60 °C catalyzed by 1.2 μM α-CT (yellow), the AuNP-Triad5 after CE (orange), the AuNP-Triad5 before CE (tangerine), the AuNP-Triad5-HSD after CE (brown) or the AuNP-CL2 after CE (blue). (**E**) Species after 48 hours hydrolysis of DEP at 60 °C catalyzed by 1.2 μM α-CT (yellow), the AuNP-Triad5 after CE (orange), the AuNP-Triad5 before CE (tangerine), the AuNP-Triad5-HSD after CE (brown) or the AuNP-CL2 after CE (blue). (**F**) Species after 48 hours hydrolysis of DBP at 60 °C catalyzed by 1.2 μM α-CT (yellow), the AuNP-Triad5 after CE (orange), the AuNP-Triad5 before CE (tangerine), the AuNP-Triad5-HSD after CE (brown) or the AuNP-CL2 after CE (blue). (**G**) Species after 72 hours hydrolysis of DEHP at 60 °C catalyzed by 1.2 μM α-CT (yellow), the AuNP-Triad5 after CE (orange), the AuNP-Triad5 before CE (tangerine), the AuNP-Triad5-HSD after CE (brown) or the AuNP-CL2 after CE (blue).

In the methanol aqueous solution, the degradation of DTP involves multiple transesterification and hydrolysis processes (Fig. 3A), and all these processes can be catalyzed by esterases. As shown in Fig. 3C and figs. S18–S22, the addition of methanol (10% v/v) makes the hydrolysis of DTP easier, and Goldenzyme shows a much higher catalytic efficiency than α-CT in both hydrolysis and transesterification. As comparisons, the AuNP-Triad5 before CE, the AuNP-Triad5-HSD after CE, and the AuNP-CL2 after CE show much lower catalytic activities for the hydrolysis and very limited activities for the transesterification of DTP (Fig. 3C).

It must be pointed out that, although DTP is more stable than *p*-NPA, there were still about 7% and 16% auto-hydrolysis after 48 hours’ incubation in the PB buffer and the buffer after CE, respectively (figs. S23, S24), indicating that DTP is not so non-activated. To further verify the “real” natural enzyme-like activity of Goldenzyme on “truly” non-activated substrates that do not show any detectable auto-hydrolysis, we then used phthalate esters (PAEs) as the substrates. Four PAEs were chosen, including dimethyl phthalate (DMP), diethyl phthalate (DEP), dibutyl phthalate (DBP), and di(2-ethylhexyl) phthalate (DEHP). PAEs are common plasticizers and durable pollutants that have caused a serious environmental problem. All these four PAEs have been listed as the priority pollutants by United State environmental protection agency (USEPA), and none of them show any sign of auto-hydrolysis in all the buffers used in this study.

Our results confirm that the longer alkyl chain of PAEs, the more difficult to be hydrolyzed (Fig. 3D–G). Remarkably, the tougher the substrate, the bigger the difference in catalytic activity between Goldenzyme and α-CT. As can be seen from Fig. 3D–G and figs. S25–S44, Goldenzyme can substantially degrade all four PAEs within 2-3 days, but α-CT only shows a much less degradation of DMP, a trace amount of degradation of DEP, and no detectable degradation of both DBP and DEHP. As dramatic comparisons, the AuNP-Triad5 before CE and the AuNP-Triad5-HSD after CE only show a trace amount of degradation of DMP, and no detectable degradation of the other three PAEs; and the AuNP-CL2 after CE shows no detectable degradation of all the four PAEs. These results unambiguously demonstrate that the high catalytic activity of Goldenzyme for the hydrolysis of non-activated esters mainly comes from the CE effects.

Taking advantages of its 20 active sites, Goldenzyme, the α-CT mimic, shows uniformly high catalytic activities for the hydrolysis of all six esters tested, including both activated and non-activated ones. Many claims of efficient activity of hydrolase-mimics in the literature based on the value of *k*_*cat*_*/k*_*uncat*_ obtained on activated esters cannot be reproduced on the hydrolysis of non-activated esters (*56*), probably due to the unreliable tiny value of *k*_*uncat*_. The consistently high catalytic efficiency of Goldenzyme for both activated and non-activated esters suggests that *k*_*cat*_, instead of *k*_*cat*_*/k*_*uncat*_, is more reliable and convincible, in particularly when a natural enzyme is used parallelly as a positive control.

### Fast-dynamic feature of Goldbody

To check whether Goldenzyme has the designed fast-dynamic feature, we used NMR spectroscopy to measure the kinetics of the hydrogen-deuterium exchange (HDX) of the backbone amide hydrogens (*67,68*). The backbone amide hydrogens are quickly exchangeable in the random-coil conformation, so all the hydrogens of the backbone amides of free Triad5 were quickly exchanged to deuterium in D_2_O, thus the peaks of the backbone amide hydrogens almost disappeared within a minute (Fig. 4A and figs. S45–S47).

**Fig. 4.**
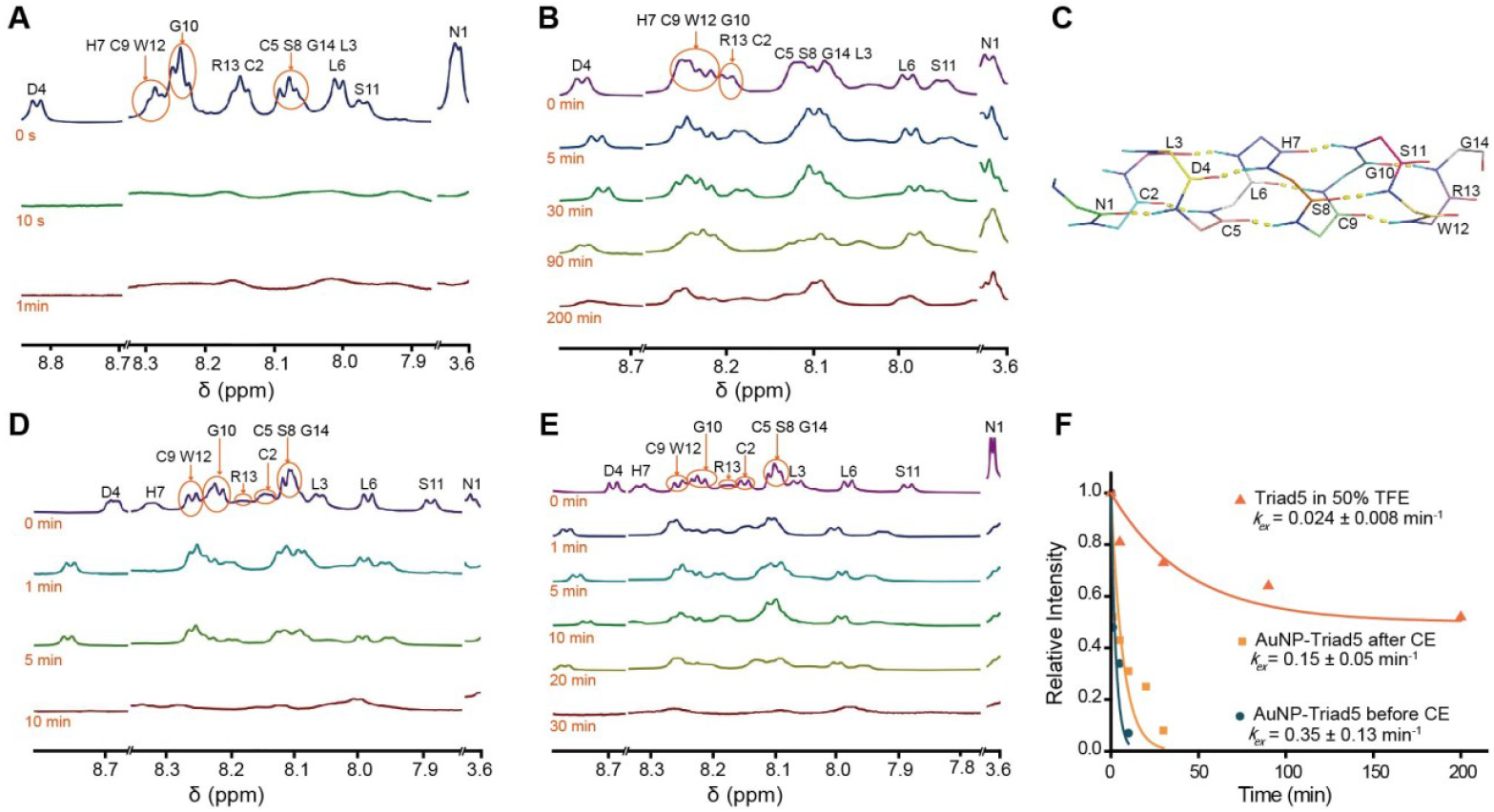
Hydrogen-deuterium exchange (HDX) kinetics of backbone amide hydrogens measured by ^1^H NMR spectra. (**A**) NMR spectra of the free Triad5 peptide. (**B**) NMR spectra of the Triad5 in 50% TFE. (**C**) Structural model of the H-bonds between residues i and i+4 in α-helical Triad5. (**D**) NMR spectra of the AuNP-Triad5 before CE. (**E**) NMR spectra of Goldenzyme (AuNP-Triad5 after CE). (**F**) HDX processes fitted to single exponential decay curves, in which the data points are the normalized peak areas of all amide hydrogens of Triad5 except for those of N1 and D4.

In 50% TFE, Triad5 folds into an α-helical conformation, thus the backbone amide hydrogen of the i+4 residue forms a hydrogen bond with the backbone carbonyl oxygen of the i residue, protecting the amide hydrogens from exchanging (Fig. 4B,C). Therefore, the backbone amide hydrogens of the Triad5 in 50% TFE show much slower average HDX rate (*k*_*ex*_) of 0.024 min^−1^ (Fig. 4B,F and fig. S48).

After conjugated onto AuNPs, the backbone amide hydrogens of Triad5 in the AuNP-Triad5 before CE show an average *k*_*ex*_ of 0.35 min^−1^ (Fig. 4D,F and fig. S49), which is much slower than that of free Triad5 peptide, indicating the shielding effect of AuNPs on the HDX.

After CE, the backbone amide hydrogens of the Triad5 of Goldenzyme (the AuNP-Triad5 after CE) show an average *k*_*ex*_ of 0.15 min^−1^ (Fig. 4E,F and fig. S50), which is more than two-fold slower than that of the AuNP-Triad5 before CE, indicating the formation of protective conformation after CE for the Triad5 of Goldenzyme, which is consistent with the CD result that the Triad5 in Goldenzyme is in an α-helical conformation (Fig. 1H). Interestingly, despite of the shielding effect of AuNPs for Goldenzyme, the average HDX rate of the backbone amide hydrogens of the Triad5 of Goldenzyme is about 6-fold faster than that of typical α-helix (the Triad5 in 50% TFE), clearly demonstrating the fast-dynamic feature of Goldenzyme.

## Discussion

Since Pauling proposed that enzymes are complementary in structure to the transition state of the substrate (*69*), preferential binding/stabilizing the transition state has been widely accepted as the mechanism of enzymatic reactions, and widely used as a general guideline for the design of enzyme-mimics, especially de novo designed enzymes. So far, many efforts have been made to mimic the catalytic triad and/or oxyanion hole of hydrolases. Yet, most of them only exhibit a limited catalytic turnover number on activated esters (*28–31*), implying that the correct combination of key residues, even in a well-defined structure that is complementary to the transition state of the substrate, does not guarantee a high catalytic activity, at least not for the hydrolysis of non-activated esters.

It was pointed out that Pauling’s complementary transition state theory cannot account for the enzymatic reactions with a very high activation free-energy (>15 kcal·mol^−1^) (*70*). Fundamentally, the biggest difference between activated and non-activated esters is not their (transition state) binding specificity and affinity with the enzymes or enzyme-mimics, but their dramatically different activation free-energy barriers. The high activation free-energy of non-activated esters might be the major reason that even those de novo enzymes, which are designed according to Pauling’s complementary transition state theory, cannot hydrolyze non-activated esters, let alone other hydrolase-mimics. Here, we developed a CE approach and designed an efficient artificial esterase that can even hydrolyze the tough non-activated esters. Apparently, the well-organized (in α-helical conformation) active center with fast-dynamic feature (about 6-fold faster than normal α-helix) of the backbone is essential for the high catalytic efficiency of Goldenzyme.

A notable difference between Goldenzyme and current de novo designed enzymes as well as those well-structured hydrolase-mimics (e.g. peptide fibers and dendrimers) is that the key catalytic residues of Goldenzyme are incorporated in flexible fragments, while those of de novo enzymes are usually fixed on more stable helices or sheets, and those of peptide fibers and dendrimers are densely packed. Placing key catalytic residues on well-organized yet flexible fragments has three potential advantages. First, the flexible fragments provide the key residues greater freedom of movement than the mere rotation of their sidechains, enabling more accurate induced-fitting with substrates (*71*). Second, as the hydrolysis of substrates contains several consecutive elementary reactions, flexible fragments allow the key residues to be transiently arranged in different correct arrangements during different hydrolysis stages (*72*). Third, the overall movement of the fragments has much greater momentum than the movement of individual sidechains or functional groups, and thus can lead to stronger and more durable collisions with the substrates to overcome the high activation free-energy barriers of non-activated esters. Given the merits of flexible fragments, we anticipate that future enzyme-mimics, especially de novo designed enzymes, would make great advances by incorporating the key residues into flexible loop fragments rather than rigid helix or sheet fragments.

Another key factor in the success of Goldenzyme is the synergy among the catalytic triad, the oxyanion hole, and the substrate-binding site, which has been unambiguously demonstrated by the results of the mutation experiments. However, Goldenzyme has only one residue (Trp) designed for the binding site, resulting in a relatively weak binding to the substrate (*p*-NPA) with a *K*_*m*_ one order of magnitude larger than those of many enzymes, leaving a large room for future improvement.

Natural proteins are always evolving slowly, but human intervention and invention can speed up this process to produce novo and better enzymes and medicines (*73–75*). An intrinsic weakness of natural proteins is their instability, which is mainly due to the instability of the protein scaffold. Together with our previous work (*76*), we have demonstrated here that AuNP could serve as an alternative stable scaffold to hold the key functional residues, and CE could be used to tune the key residues into the right conformation. In fact, CE could also be applied to NPs other than AuNPs (*52,77,78*). By replacing the instable scaffolds of natural proteins with stable inorganic NPs, NP-based protein mimics via CE would surpass natural proteins in many aspects (*76,79*) and replace them in future industrial and biomedical applications.

## Materials and methods

### Materials

Chloroauric acid, sodium tetrahydroborate, and trisodium citrate dehydrate were purchased from Sinopharm Chemical Reagent (China). Peptides (95% purity) were synthesized by GL Biochem. Ltd. (China). 2,2,2-trifluoroethanol (TFE, 99.5% purity) were bought from Aladdin Reagent (China). α-chymotrypsin (α-CT, type I-S, essentially salt-free, lyophilized powder), *p*-nitrophenol (*p*-NP, 99% purity) and *p*-nitrophenyl acetate (*p*-NPA, 99% purity) were purchased from Sigma-Aldrich (USA). Diethyl terephthalate (DTP, 98% purity), I_2_ (99.8% purity), Na_2_S_2_O_3_ (99% purity), and KI (99% purity) were purchased from Shanghai Macklin Biochemical Co., Ltd. (China). Dimethyl phthalate (DMP, 99% purity), diethyl phthalate (DEP, 99% purity), dibutyl phthalate (DBP, 99% purity), and di(2-ethylhexyl) phthalate (DEHP, 99% purity), D_2_O (99.5% purity), deuterated DMSO-d6 (99.8% purity), deuterated formic acid-d2 (98% purity), deuterated TFE-d3 (98% purity) were purchased from Adamas Reagent (China). All other reagents are of analytical grade from commercial sources. Ultrapure water (18.2 MΩ·cm) was used.

### Synthesis and characterization of AuNPs

Citrate-capped AuNPs (∼3.6 nm) were synthesized and quantified as previously reported (*42–48*). The concentration of AuNPs was calculated from the absorbance at 450 nm using an ε_450 nm_ of 2.768 × 10^6^ M^−1^·cm^−1^ (*80*). The as-prepared AuNPs were stored at room temperature in the dark, and filtered with a 0.22 μm filter before use.

### Peptide conjugation and CE to prepare Goldenzyme and its mutants

The concentration of peptides was determined by their absorbance at 280 nm using estimated extinction coefficients (http://web.expasy.org/protparam/)(see **table S1**). In a typical procedure for synthesizing AuNP-peptide conjugates, 10 μL of 0.2 M trisodium citrate solution was added to 6 mL of the filtered AuNPs solution under gentle stirring, which improved the stability of the solution. Next, 2 mL of the peptide solution and 50 μL 0.2 M sodium hydroxide were added dropwise into the above AuNP solution. The mixture was stirred for 1 hour at room temperature to complete the conjugation. The peptide density on AuNPs was easily regulated by adjusting the concentration of the peptide in the above procedure.

The efficiency of conjugating peptides to AuNPs was determined by comparing fluorescence spectra of the peptide before conjugation and the free peptide after conjugation. The free peptide after conjugation was the peptide remained in the filtrate after ultrafiltration (Millipore Amicon Ultra-15 filter, MW cut-off: 30k, 1500g, 15 min, 25 °C). Fluorescence spectra were recorded at 25 °C using a FS5 fluorescence spectrometer (Edinburgh Instruments, UK) with an excitation wavelength of 280 nm and a slit of 5 nm.

For CE treatment, TFE was added to the solution of AuNP-peptide conjugates (to a final concentration of 50% (v/v) TFE), and the solution was incubated at 25 °C, 37 °C, 45 °C, or 55 °C for 12, 24, or 48 hours in an incubator (Yiheng Scientific Instruments Co., Ltd., China). After incubation, the solution was frozen immediately with liquid nitrogen, followed by lyophilization in a Christ Alpha 1-2 LDplus freeze-dryer (Germany) for 72 hours. The obtained powders of AuNP-peptide conjugates are soluble in water.

### Characterization of AuNPs, Goldenzyme and its mutants

The size and morphology of AuNPs, Goldenzyme and its mutants were characterized by high-resolution transmission electron microscope (HRTEM, JEM-2100F, JEOL, Japan) and absorption spectra (U-3010, Hitachi, Japan). The hydrodynamic diameters of AuNPs, Goldenzyme and its mutants were measured by the dynamic light scattering (DLS) method using a ZS90 Zetasizer (Malvern, UK) at 25 °C.

To check whether TFE has been completely removed by lyophilization, the Goldenzyme powder was dissolved in D_2_O to a concentration of 1 μM, and then detected by ^19^F NMR on a Bruker Advance III 600 MHz NMR spectrometer. Pure TFE dissolved in D_2_O to a concentration of 100 mM was detected as the standard.

### Circular dichroism (CD) spectroscopy

The far-UV CD spectra were recorded at 25 °C in a 0.1 cm path-length quartz cuvette on JASCO J-1500 (Japan) (1 nm bandwidth, 1 s response, 0.5 nm data pitch, and at 100 nm/min scan rate). The concentration of free peptides and the peptides in AuNP-peptide conjugates was 0.1 mg·mL^−1^. To reduce noise, the final CD spectra were averages of 20 scans.

### Determination of the catalytic efficiency for the hydrolysis of p-NPA

To eliminate the influence of the high ionic strength and all solutes in the solution of Goldenzyme, we ultrafiltrated the AuNP-peptide (before CE) solution and used the filtrate as the buffer before CE, which contained the same concentration of all solutes as in the AuNP-peptide before CE solution. The buffer before CE was used as the background for the data of the AuNP-peptide before CE. The buffer after CE was prepared by treating the buffer before CE with TFE in the same way as the CE treatment of Goldenzyme, so the buffer after CE contained the same concentration of all solutes including trace TFE as in the Goldenzyme solution. And the buffer after CE was used as the background for the data of Goldenzyme and its mutants, thus, the contribution of ions and trace TFE had been eliminated.

The hydrolysis of *p*-NPA by the as-prepared citrate-capped AuNPs without peptide-conjugation in both the buffer before CE and the buffer after CE were also examined.

The catalytic activities of AuNP-CL2 before CE and AuNP-CL2 after CE for the hydrolysis of *p*-NPA were measured similarly, and the obtained data were used as the more accurate blanks for those of the AuNP-peptide conjugates before CE and after CE (including Goldenzyme), respectively.

For the hydrolysis of *p*-NPA, the *p*-NPA stock solution was freshly prepared by dissolving *p*-NPA in acetonitrile, and then diluting in water (the final acetonitrile content in the hydrolysis mixture was 2% (v/v)). The catalytic reaction was carried out at 25 °C in 40 mM phosphate buffer (PB, Na_2_HPO_4_-NaH_2_PO_4_, pH = 7.0), using a typical final catalyst (Goldenzyme, the mutants and blanks) concentration of 0.5 μM. The hydrolysis was initiated by automated mixing the stock solutions of the buffer, the catalyst solution and the *p*-NPA stock solution in a volume ratio of 0.5 : 5.5 : 4.0 using an SFM-3000/S stopped-flow instrument (Bio-logic, France).

The yield of the product, *p*-NP, was monitored by recording the absorbance increase at 400 nm with a MOS-500 spectrophotometer (Bio-logic, France) using a flow cell with a pathlength of 0.15 cm, and quantified using an experimentally determined molar extinction coefficient ε_400 nm_ of 10.85 mM^−1^·cm^−1^.

The initial catalytic rate (*v*_*0*_) of the hydrolysis of *p*-NPA at different substrate concentrations ([*S*], ranging from 0.04 to 3.6 mM) was calculated as the initial slope of the monitored hydrolysis process. After subtraction of the initial catalytic rate of the corresponding blanks, the *v*_*0*_ values of Goldenzyme and its mutants versus [*S*] were fitted to the Michaelis–Menten equation (1):

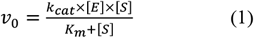

where *k*_*cat*_ is the catalytic turnover number, *K*_*m*_ is the substrate binding constant, and [*E*] and [*S*] are the concentrations of the enzyme and the substrate, respectively. Reported results are the averages of at least three independent measurements.

For a more reliable comparison between Goldenzyme and α-CT, desalted α-CT was dissolved in the buffer after CE, so that both Goldenzyme and α-CT were assayed in the same buffer. As the activity of α-CT is much lower than that of Goldenzyme, the concentration of α-CT (10 μM) that we used for the Michaelis–Menten equation fitting was 20 times that of Goldenzyme (0.5 μM).

To enable comparison with values reported in the literature, the catalytic activities of α-CT and PLE were also assayed in PB buffer.

### Thermal stability assay

The Goldenzyme and α-CT solutions were sealed in tubes and boiled in a water bath for 1 hour, then naturally cooled to 25 °C. After that, the catalytic activity assays and CD spectroscopic measurements were carried out as described above.

### Degradation of DTP

DTP was dissolved in acetonitrile or methanol, and diluted with water (the final acetonitrile or methanol content in the hydrolysis mixture was 10% (v/v)). The catalytic activity assays were carried out in 160 mM PB (pH = 8.0) containing 5 mM DTP. Reactions were initiated by mixing the corresponding buffer stock solution, DTP, and the catalyst (Goldenzyme, α-CT (dissolved in the buffer after CE), or the controls. Their final concentration was 0.8 μM). The mixtures were incubated at 37 °C in a thermomixer (200 rpm, RCTS100, Rui Cheng, China). After incubation, the mixtures were ultrafiltrated (Millipore Amicon Ultra-15 filter, MW cut-off: 10k, 5000g, 30 min, 25 °C) to separate the hydrolyzed products from enzymes.

The hydrolyzed products were analyzed using an ACQUITY UPLC H-Class series liquid chromatography-mass spectrometer (LC-MS, Waters, USA). Samples (5 μL) were loaded onto a reversed phase column HSS-C18 (ACQUITY UPLC C18: particle size, 1.8 µm; pore size, 95 Å; length, 50 mm; inside diameter, 2.1 mm) and eluted using an acetonitrile/water (5/95, v/v) mobile phase at a flow rate of 0.4 mL·min^−1^, and monitored at a wavelength of 241 nm. Mass spectra were acquired using an ion trap-type instrument (Waters SQ Detector, USA) equipped with an electrospray ion source.

### Recyclability of Goldenzyme after the degradation of DTP

After 48 hours of the above degradation of DTP, the solution was extracted twice with ethyl acetate to remove the products and unreacted substrate. The extracted mixture was detected by the LC-MS as described above, and the remaining Goldenzyme solution was used for the next cycle of the degradation of fresh DTP substrate. The process was repeated 4 times.

### Degradation of DMP, DEP, DBP, and DEHP

DMP, DEP, DBP and DEHP were dissolved in acetonitrile and diluted with water (the final acetonitrile content in the hydrolysis mixture was 10% (v/v)). The degradation assays were carried out in 160 mM PB (pH = 8.0) containing 5 mM DMP, DEP, DBP or DEHP. Similar to the degradation of DTP, the reactions were initiated by mixing the corresponding buffer stock solution, the substrate and the catalyst (Goldenzyme, α-CT (dissolved in the buffer after CE), or the controls. Their final concentration was 1.2 μM). The mixtures were incubated at 60 °C in the thermomixer (200 rpm). After incubation for 48 or 72 hours, the mixtures were separated and analyzed as described above.

### Hydrogen–deuterium exchange (HDX) kinetics detected by NMR

All NMR spectra were acquired on a Bruker 600 MHz spectrometer. Chemical shifts are reported in ppm. ^1^H NMR signals were assigned from the 1H-1H COSY.

For the AuNP-Triad5 before (control) and after (Goldenzyme) CE, the HDX was initiated by dissolving 5 mg of lyophilized samples in 0.55 mL D_2_O. After certain periods of time for HDX, 4 µL of ice-cold formic acid-d2 was added to quench the HDX (final pH = 3.0) (*68*), followed by the addition of 50 µL of the KI/I_2_ etching solution to dissolve the AuNPs (*81*), resulting in a yellow solution in a few seconds. Then, 100 µL of Na_2_S_2_O_3_ was added to eliminate excess I_2_, resulting in a colorless, transparent solution in a few seconds. The solution was immediately immersed in liquid nitrogen, followed by lyophilization. The lyophilized powders were re-dissolved in 0.55 mL of DMSO-d6 for NMR analysis.

As 5 mg of Goldenzyme dissolving in 0.55 mL D_2_O resulted in a solution with a pH of 11.7, to carry out the HDX detection of free Triad5 under the same condition, 5 mg of pure Triad5 powder was dissolved in 0.5 mL of the buffer after CE (pH = 11.7), and then lyophilized to remove all H_2_O. Subsequently, the lyophilized Triad5 was dissolved in 0.55 mL of D_2_O to initiate the HDX. After certain periods of time, the HDX was quenched by adding 4 µL of ice-cold formic acid-d2, and the sample was immediately immersed in liquid nitrogen and then lyophilized. The lyophilized powders were re-dissolved in 0.55 mL of DMSO-d6 for NMR analysis.

For the HDX detection of the Triad5 in 50% TFE, 5 mg of Triad5 was dissolved in 0.5 mL of the buffer after CE (pH = 11.7), then 0.5 mL of TFE was added. The mixture was incubated for 5 hours to complete the induction of α-helix. Subsequently, the HDX was initiated by adding 1 mL of the TFE-d3 (50% v/v) D_2_O solution. After certain periods of time, the HDX was quenched by adding 4 µL of ice-cold formic acid-d2, and the sample was immediately immersed in liquid nitrogen and then lyophilized. The lyophilized powders were re-dissolved in 0.55 mL of DMSO-d6 for NMR analysis.

The NMR peaks of the backbone amide hydrogens of Triad5 were quantified using the peaks of the stable hydrogens of the methyl groups of the leucine residue (L3) of Triad5 as the reference, and normalized using the corresponding value at time zero as the unit. The HDX rates were obtained by plotting the normalized backbone amide hydrogens *vs* time and fitting them to a single exponential decay function, as described by the equation (2):

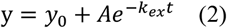

where, *y* is the normalized backbone amide hydrogens, *t* is the time of HDX, *k*_*ex*_ is the HDX rate. For the AuNP-Triad5 before CE and Goldenzyme, *y*_*0*_ = 0, *A* = 1; for Triad5 in 50% TFE, *y*_*0*_ = 0.5, *A* = 0.5, because the HDX was carried out in 50% deuterated solvents. As the HDX of free Triad5 was too fast, no enough data were obtained for reliable fitting.

Because the peak of the backbone amide hydrogen of N1 overlapped with the peak of other type of hydrogen, the total peak areas of the backbone amide hydrogens of all residues, except for N1, were used to calculate the average *k*_*ex*_.

## Acknowledgments

We would like to thank Professor Yuanfang Liu for his long-standing support and encouragement. This work was supported by the National Natural Science Foundation of China (Nos. 32371318, 22071145, 31871007, 32071404, and 31771105) and the National Key Research and Development Plan of China (No. 2016YFA0201602).

## Author contributions

Conceptualization: AC,

HW Methodology: AC, YW, YC, HW

Investigation: YW, YYL, YC, YyC, TG, JD, TZ

Visualization: AC, YW, YYL, HW

Funding acquisition: AC, HW

Project administration: AC, HW

Supervision: AC, HW

Writing – original draft: AC, YW, HW

Writing – review & editing: AC, YW, HW, YYL, YC, YyC, TG, JD, TZ

## Supplementary Materials

Figs. S1 to S50

Tables S1 to S3

