## Supplemental Figs.1-50, Table S1-S3 for "An efficient artificial esterase with a dynamic conformation via conformational engineering"

**The PDF file includes:**

Figs. S1 to S50
Tables S1 to S3

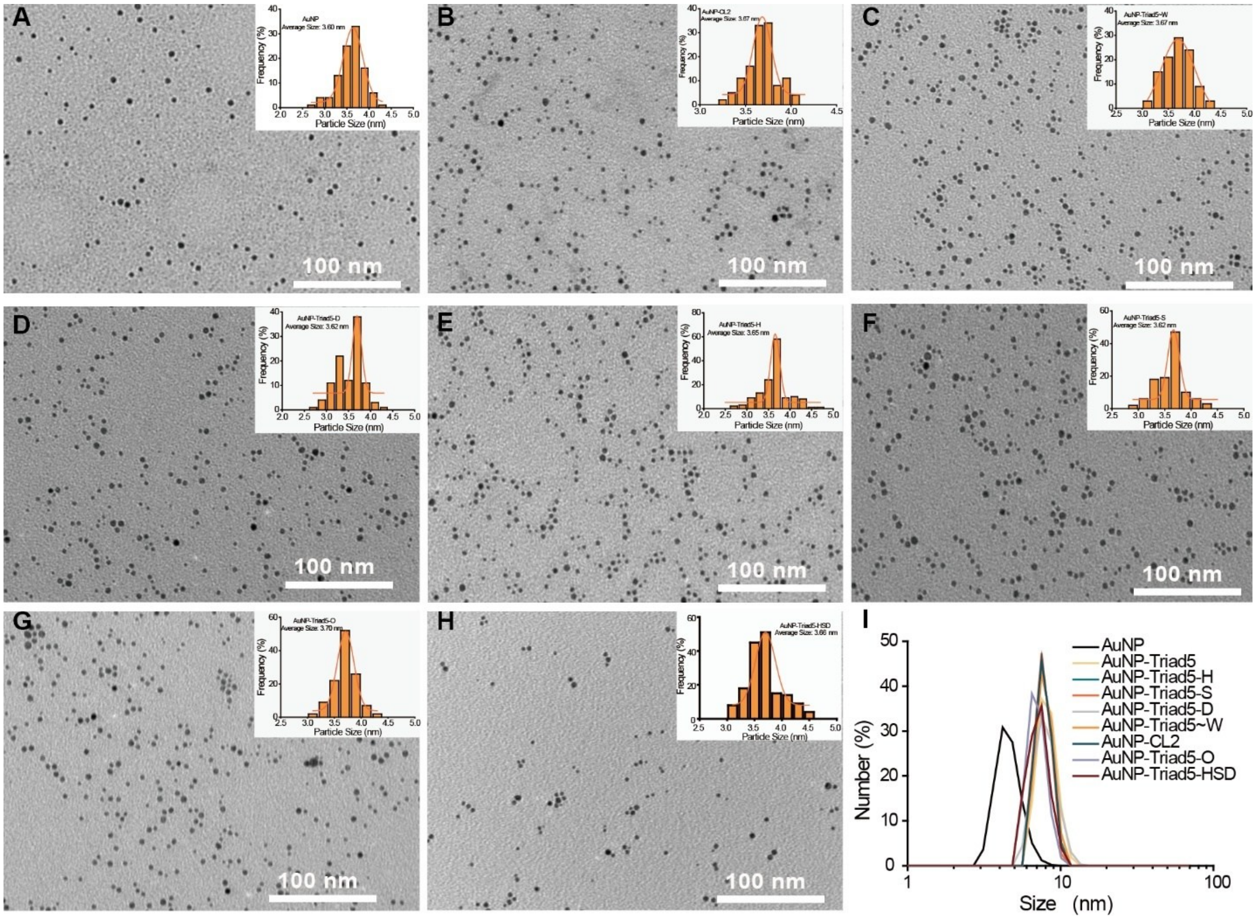


**Fig. S1. Characterization of AuNPs and the AuNP-peptide conjugates. (A-H)** TEM image and size distribution of (A) AuNPs, (B) AuNP-CL2, (C) AuNP-Triad5~W, (D) AuNP-Triad5-D, (E) AuNP-Triad5-H, (F) AuNP-Triad5-S, (G) AuNP-Triad5-O, and (H) AuNP-Triad5-HSD. **(I)** Hydrodynamic diameters (DLS results) of AuNPs and the AuNP-peptide conjugates.

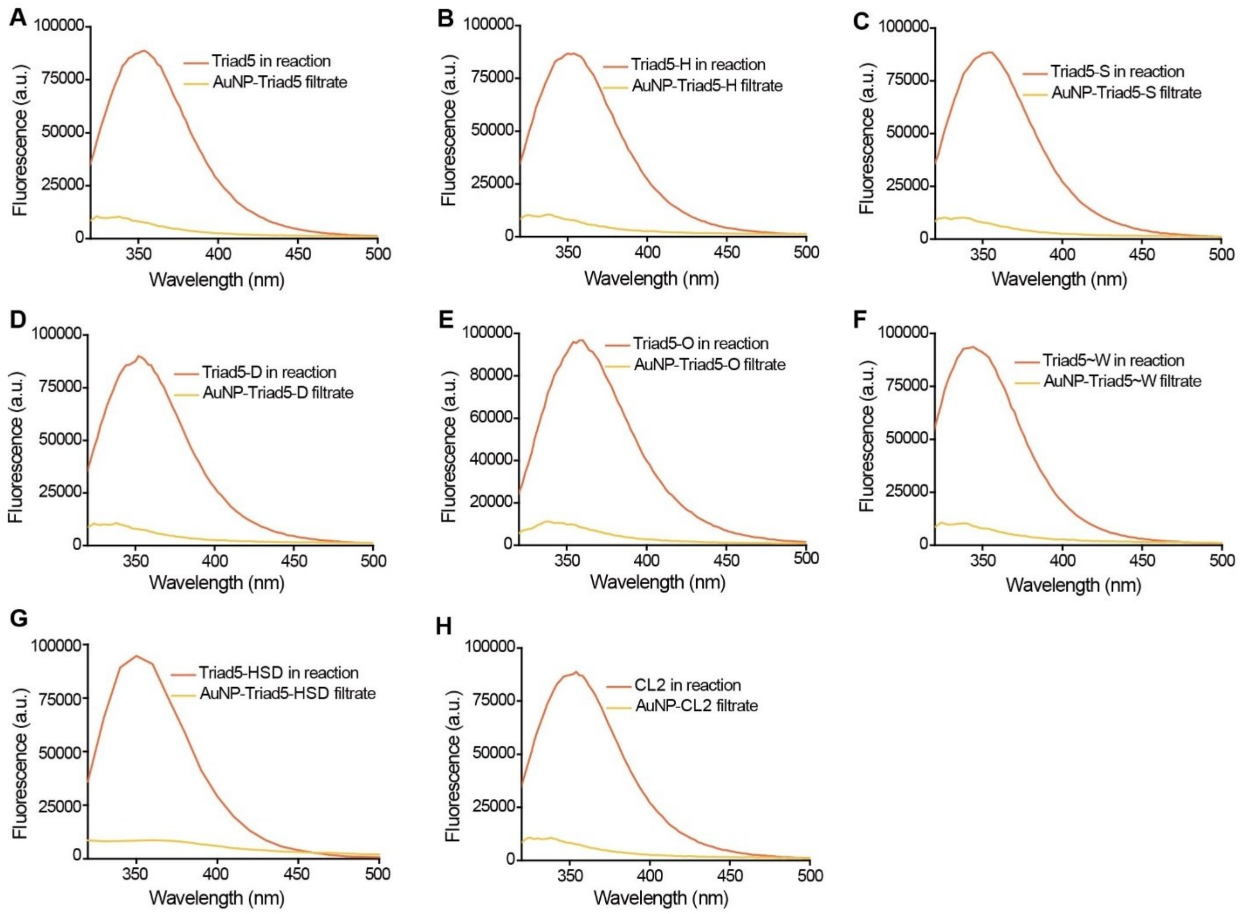


**Fig. S2. Conjugation efficiency of peptides to AuNPs as shown by the fluorescence spectra (excited at 280 nm) of the total peptide before conjugation (tangerine) and the free peptide after conjugation (i.e. free peptide in the filtrate after ultrafiltration) (yellow). (A)** AuNP-Triad5. **(B)** AuNP-Triad5-H. **(C)** AuNP-Triad5-S. **(D)** AuNP-Triad5-D. **(E)** AuNP-Triad5-O. **(F)** AuNP-Triad5~W. **(G)** AuNP-Triad5-HSD. **(H)** AuNP-CL2.

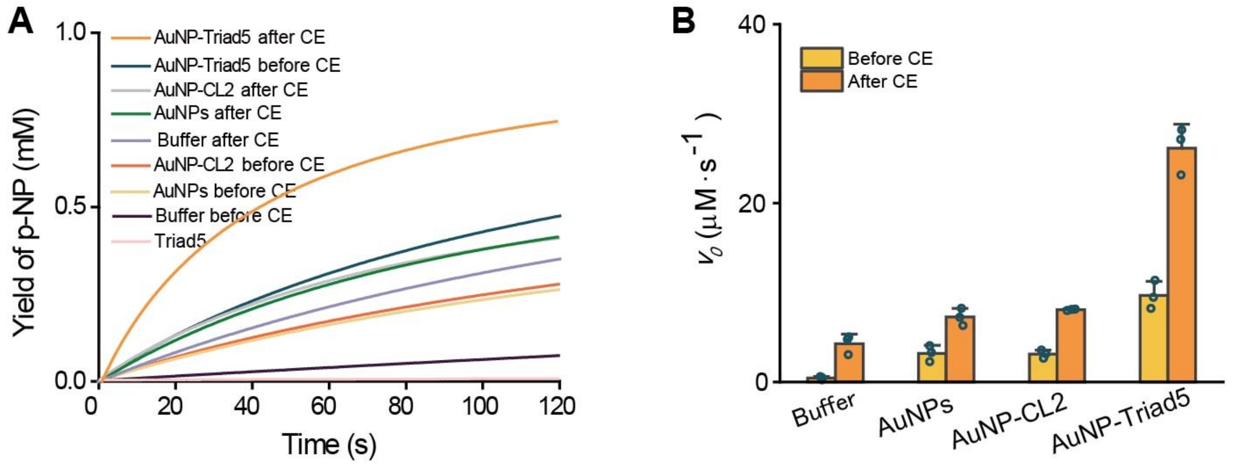


**Fig. S3. Hydrolysis process (A) and catalytic activity (B) of Goldenzyme (AuNP-Triad5 after CE), the controls and backgrounds.** Conditions for the hydrolysis of *p*-NPA: [AuNPs or AuNP-peptide conjugates] = 0.5 μM, [Triad5] = 10 μM, [S] = 1 mM, [acetonitrile] = 2% (v/v), [PB] = 40 mM, pH = 7.0, $T$ = 25 ℃.

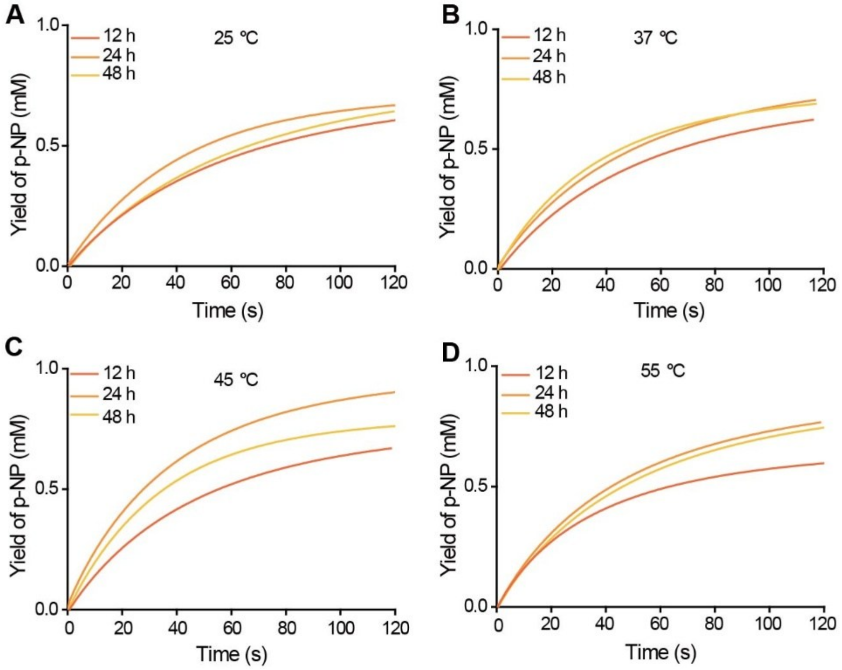


**Fig. S4. Optimization of CE induction conditions based on the hydrolysis of *p*-NPA (1 mM) catalyzed by the AuNP-Triad5 conjugate being induced by TFE at different temperatures.** **(A)** 25 °C. **(B)** 37 °C. **(C)** 45 °C. **(D)** 55 °C. Conditions for the hydrolysis: [AuNP-Triad5] = 0.5 μM, [S] = 1 mM, [acetonitrile] = 2% (v/v), [PB] = 40 mM, pH = 7.0, $T$ = 25 °C.

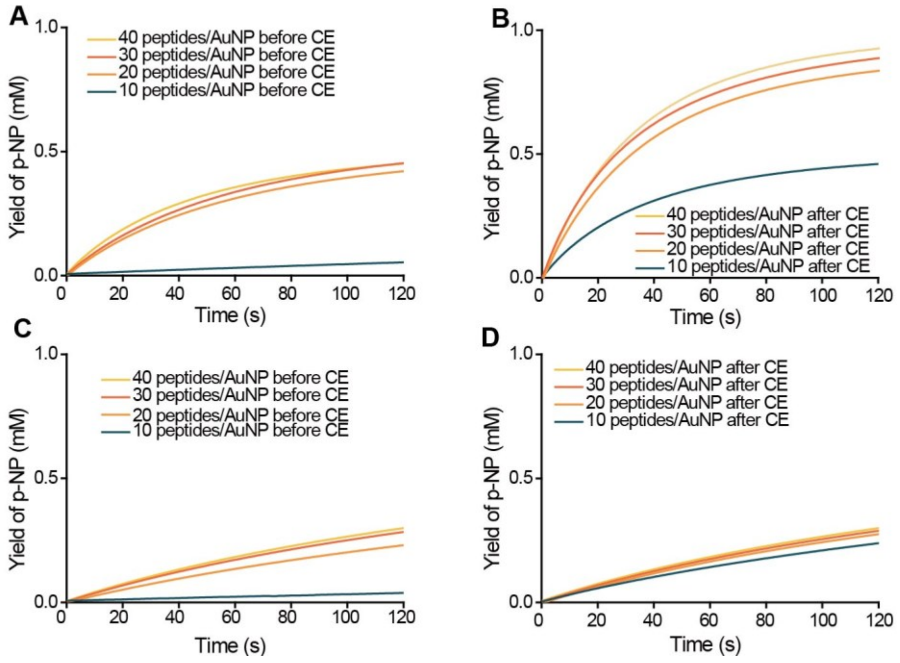


**Fig. S5. Optimization of peptide density based on the hydrolysis of *p*-NPA (1 mM) catalyzed by the AuNP-Triad5 conjugates of different peptide densities before (A) and after (B) CE, and by the AuNP-CL2 conjugates of different peptide densities before (C) and after (D) CE.** Conditions for the hydrolysis: [AuNP-peptide] = 0.5 μM, [S] = 1 mM, [acetonitrile] = 2% (v/v), [PB] = 40 mM, pH = 7.0, $T$ = 25 °C.

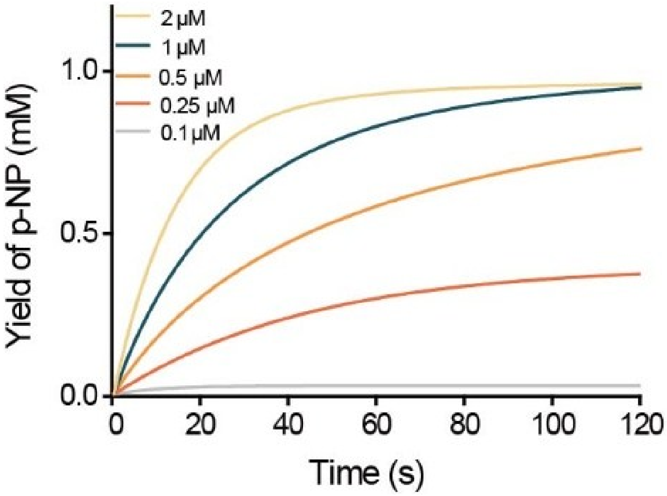


**Fig. S6. Hydrolysis processes of *p*-NPA (1 mM) catalyzed by different concentrations of Goldenzyme.** Conditions for the hydrolysis: [acetonitrile] = 2% (v/v), [PB] = 40 mM, pH = 7.0, $T$ = 25 °C.

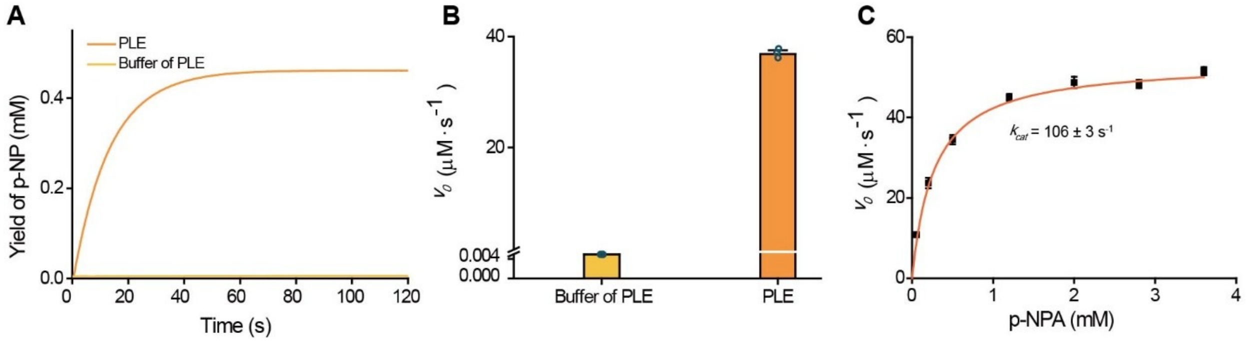


**Fig. S7. Hydrolysis of *p*-NPA (1 mM) catalyzed by porcine liver esterase (PLE, 0.5 μM). (A)** Hydrolysis process of PLE and its buffer. **(B)** Catalytic activity of PLE and its buffer. **(C)** Catalytic activity fitted to the Michaelis–Menten equation. Conditions for the hydrolysis: [E] = 0.5 μM, [acetonitrile] = 2% (v/v), [PB] = 10 mM, pH = 7.0, $T$ = 25 °C.

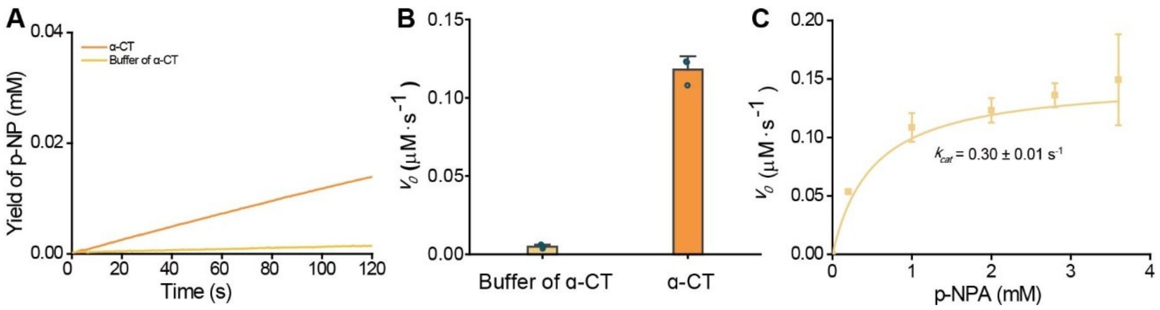


**Fig. S8. Hydrolysis of *p*-NPA (1 mM) catalyzed by α-CT (0.5 μM). (A)** Hydrolysis process of α-CT and its buffer. **(B)** Catalytic activity of α-CT and its buffer. **(C)** Catalytic activity fitted to the Michaelis–Menten equation. Conditions for the hydrolysis: [E] = 0.5 μM, [acetonitrile] = 2% (v/v), [PB] = 10 mM, pH = 7.0, $T$ = 25 °C.

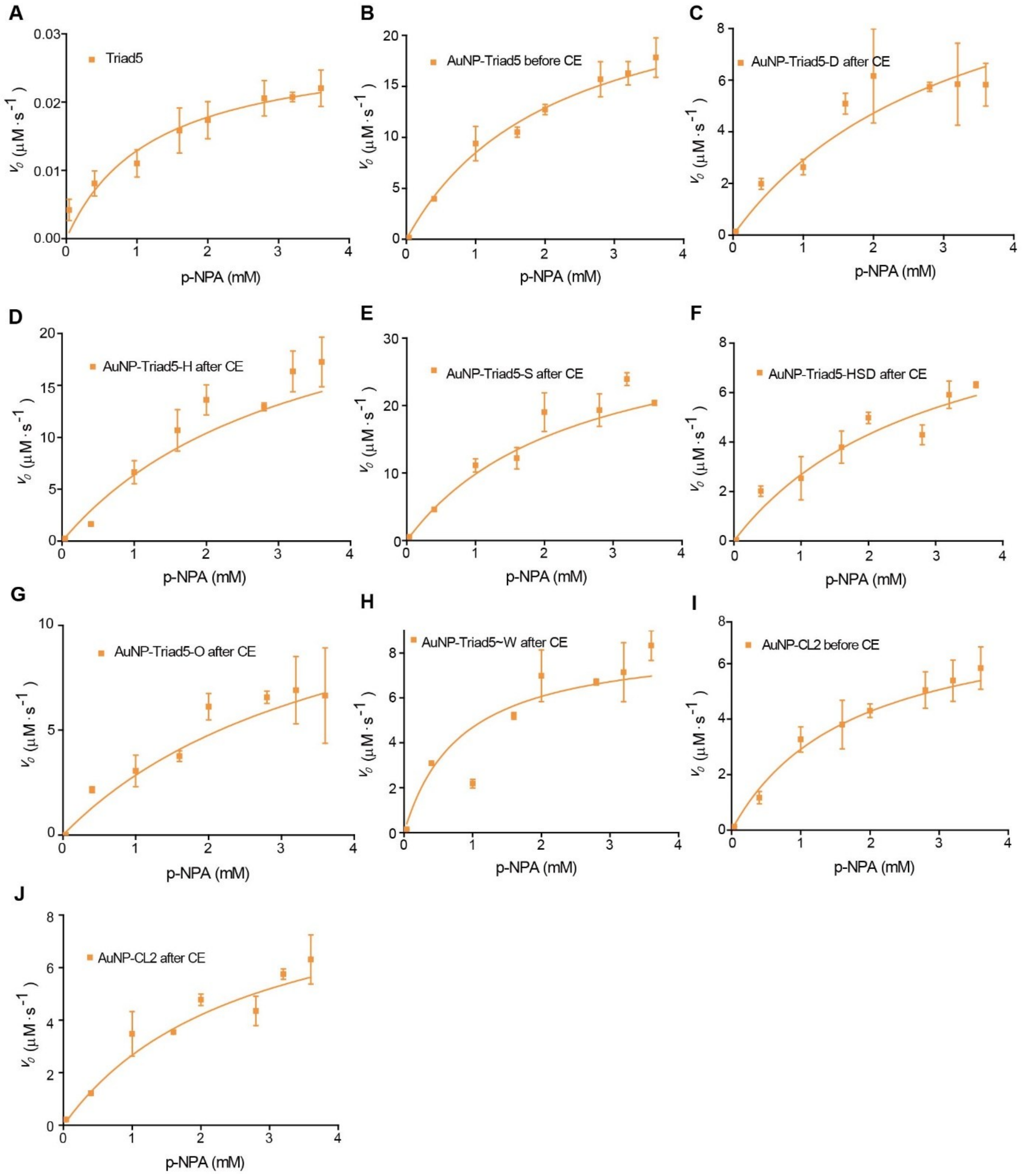


**Fig. S9. Catalytic activity of AuNP-peptide conjugates for the hydrolysis of *p*-NPA was fitted to the Michaelis–Menten equation. (A)** Triad5. **(B)** AuNP-Triad5 before CE. **(C)** AuNP-Triad5-D after CE. **(D)** AuNP-Triad5-H after CE. **(E)** AuNP-Triad5-S after CE. **(F)** AuNP-Triad5-HSD after CE. **(G)** AuNP-Triad5-O after CE. **(H)** AuNP-Triad5~W after CE. **(I)** AuNP-CL2 before CE. **(J)** AuNP-CL2 after CE. Conditions for the hydrolysis: [AuNP-peptide] = 0.5 μM, [Peptide] = 10 μM, [acetonitrile] = 2% (v/v), [PB] = 40 mM, pH = 7.0, $T$ = 25 °C. Error bars indicate the SDs of triplicate experiments.

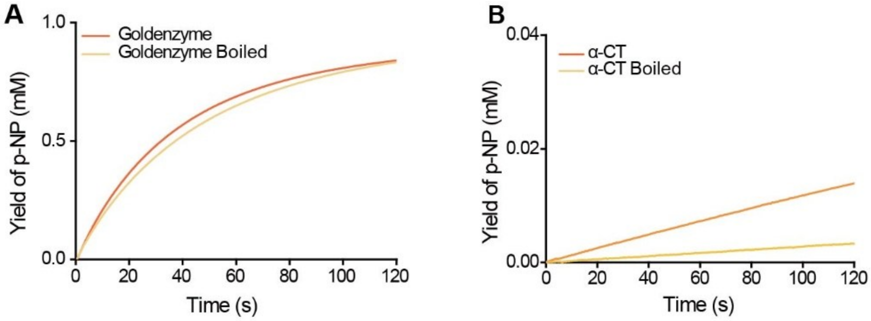


**Fig. S10. Thermal stability shown by the hydrolysis of *p*-NPA catalyzed by (A) Goldenzyme, and (B) α-CT.** Tangerine lines: intact enzymes. Yellow lines: enzymes were boiled for 1 hour followed by natural cooling to 25 °C. Conditions for the hydrolysis: [Goldenzyme] = 0.5 μM, [α-CT] = 0.5 μM, [S] = 1 mM, [acetonitrile] = 2% (v/v), [PB] = 40 mM, pH = 7.0, $T$ = 25 °C.

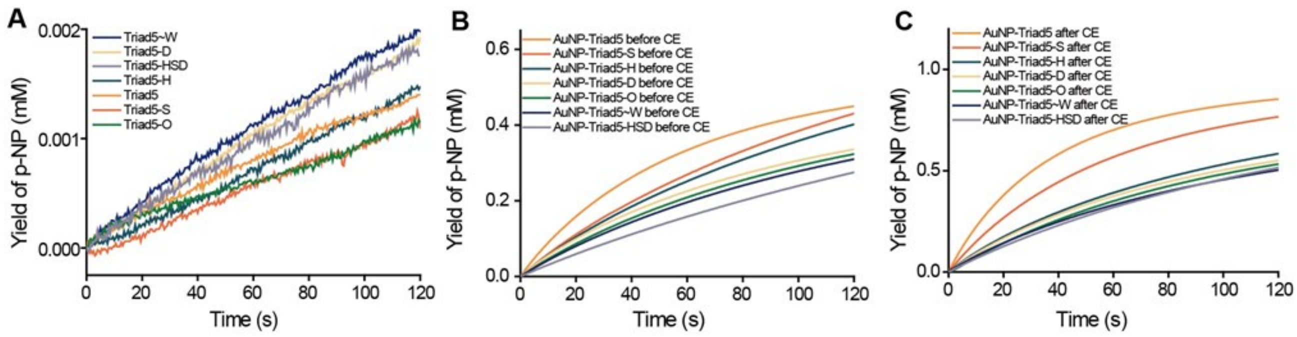


**Fig. S11. Hydrolysis of *p*-NPA catalyzed by the AuNP-peptide conjugates before and after CE, as well as the corresponding free peptides. (A)** Hydrolysis processes catalyzed by free peptides. **(B)** Hydrolysis processes catalyzed by the AuNP-peptide conjugates before CE. **(C)** Hydrolysis processes catalyzed by the AuNP-peptide conjugates after CE. Conditions for the hydrolysis: [AuNP-peptide] = 0.5 μM, [Peptide] = 10 μM, [S] = 1 mM, [acetonitrile] = 2% (v/v), [PB] = 40 mM, pH = 7.0, $T$ = 25 °C.

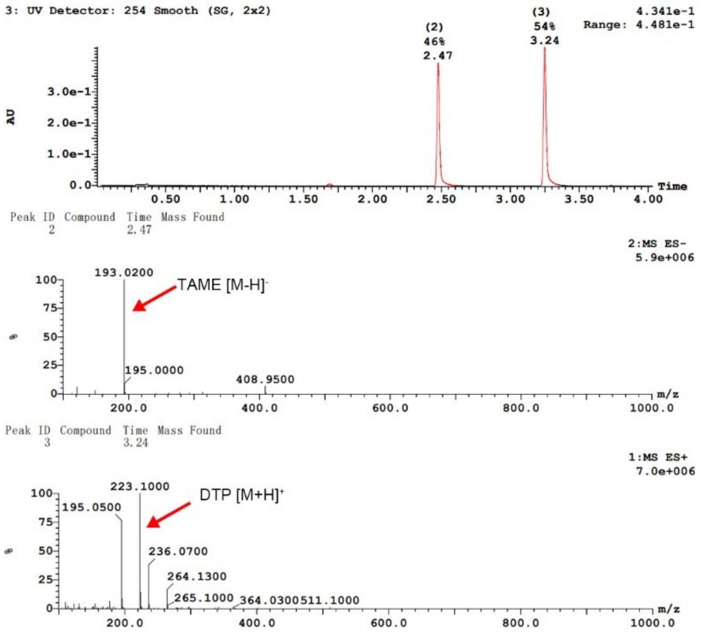


**Fig. S12. LC-MS analysis of the hydrolysis products of DTP catalyzed by α-CT (dissolved in the buffer after CE) for 48 hours.** Conditions for the hydrolysis: [E] = 0.8 μM, [S] = 5 mM, [acetonitrile] = 10% (v/v), [PB] = 160 mM, pH = 8.0, *T* = 37 °C.

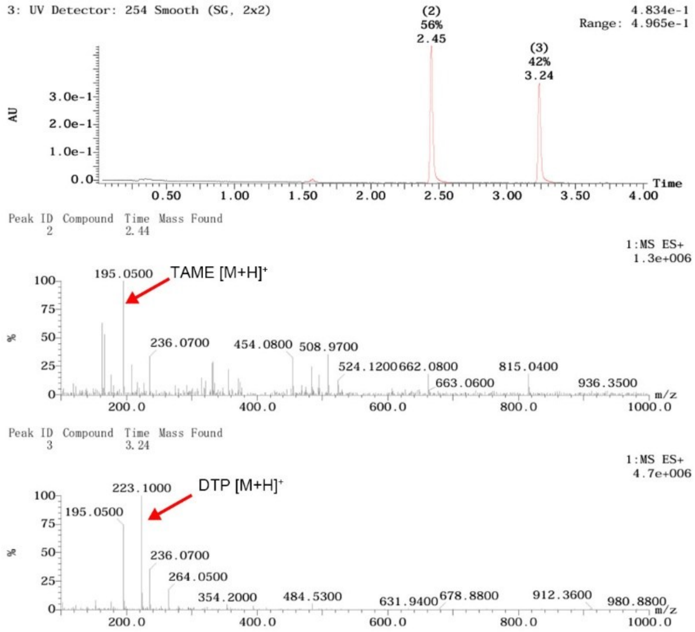


**Fig. S13. LC-MS analysis of the hydrolysis products of DTP catalyzed by Goldenzyme for 48 hours.** Conditions for the hydrolysis: [E] = 0.8 μM, [S] = 5 mM, [acetonitrile] = 10% (v/v), [PB] =160 mM PB, pH = 8.0, $T$ = 37 °C.

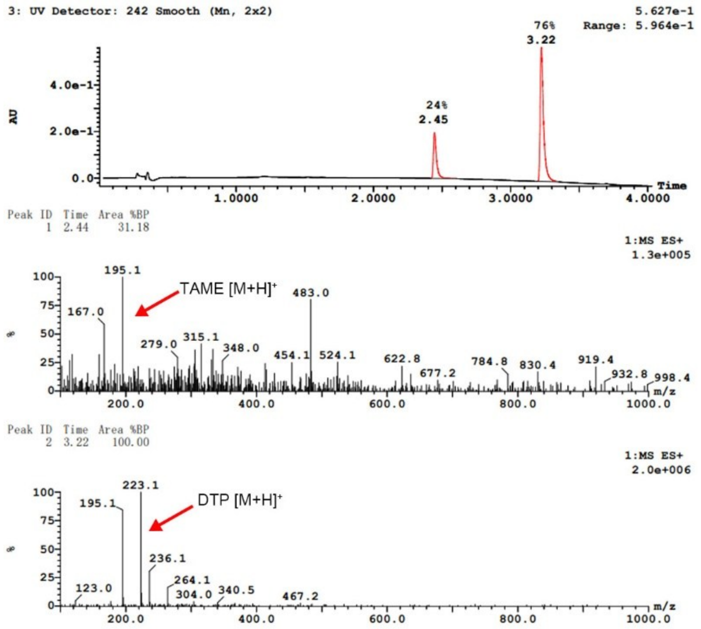


**Fig. S14. LC-MS analysis of the hydrolysis products of DTP catalyzed by the AuNP-Triad5 before CE for 48 hours.** Conditions for the hydrolysis: [E] = 0.8 μM, [S] = 5 mM, [acetonitrile] = 10% (v/v), [PB] =160 mM PB, pH = 8.0, *T* = 37 °C.

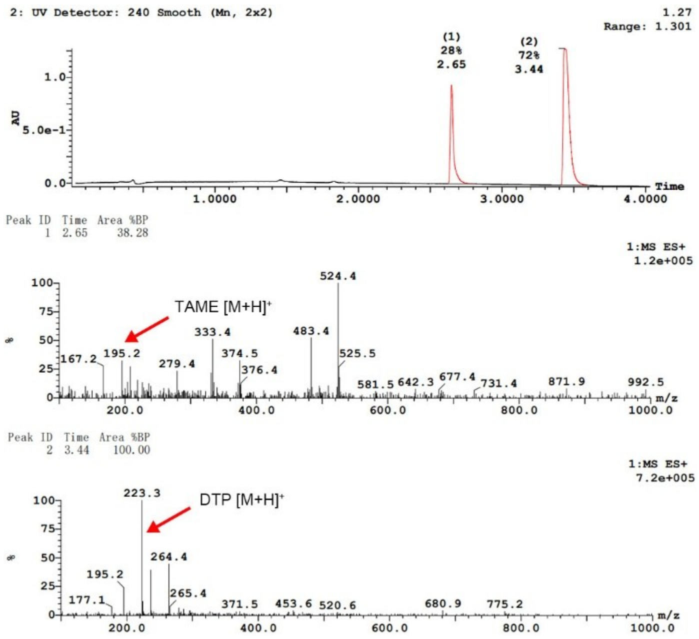


**Fig. S15. LC-MS analysis of the hydrolysis products of DTP catalyzed by the AuNP-Triad5-HSD after CE for 48 hours.** Conditions for the hydrolysis: [E] = 0.8 μM, [S] = 5 mM, [acetonitrile] = 10% (v/v), [PB] =160 mM PB, pH = 8.0, $T$ = 37 °C.

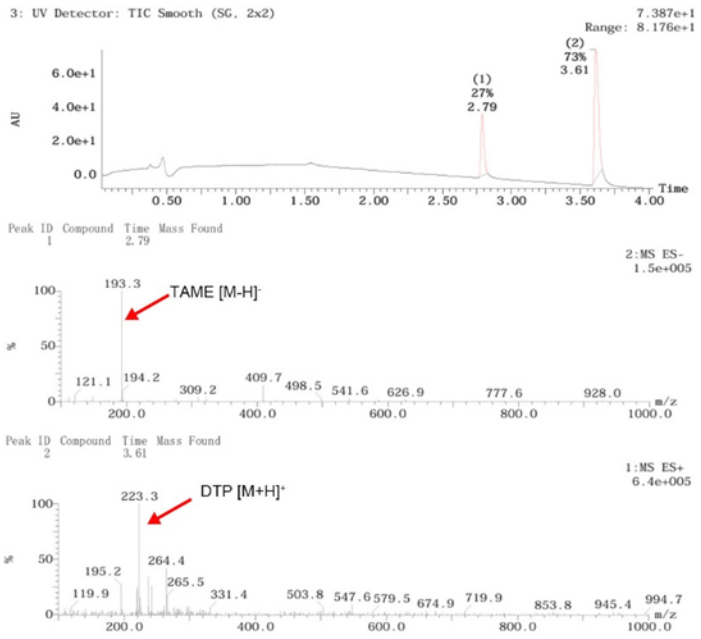


**Fig. S16. LC-MS analysis of the hydrolysis products of DTP catalyzed by the AuNP-CL2 after CE for 48 hours.** Conditions for the hydrolysis: [E] = 0.8 μM, [S] = 5 mM, [acetonitrile] = 10% (v/v), [PB] =160 mM PB, pH = 8.0, $T$ = 37 ℃.

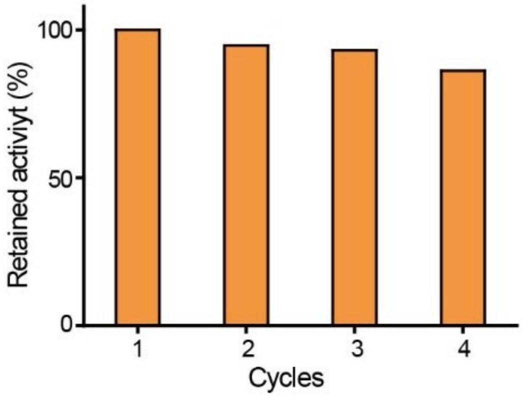


**Fig. S17. Catalytic activity of Goldenzyme after different recovery cycles (48 hours of hydrolysis for each cycle).** Conditions for the hydrolysis: [Goldenzyme] = 0.8 μM, [DTP] = 5 mM, [acetonitrile] = 10% (v/v), [PB] = 160 mM, pH = 8.0, $T$ = 37 °C.

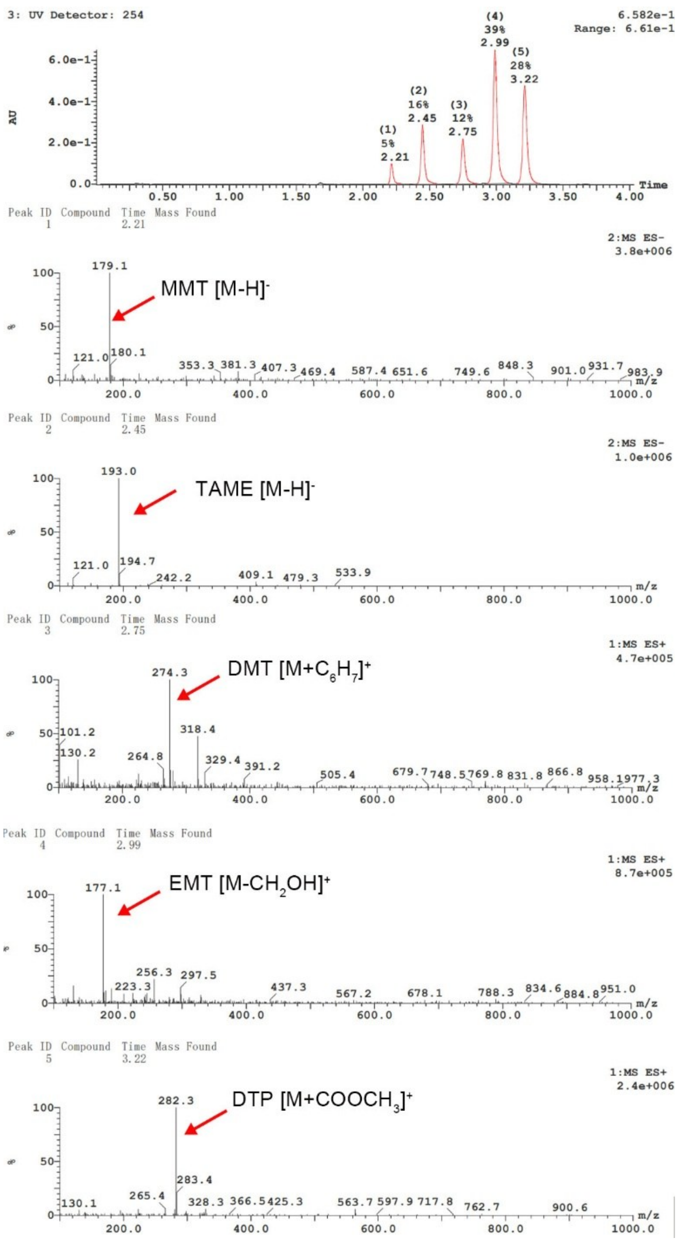


**Fig. S18. LC-MS analysis of the hydrolysis products of DTP catalyzed by α-CT (dissolved in the buffer after CE) for 24 hours.** Conditions for the hydrolysis: [E] = 0.8 μM, [S] = 5 mM, [methanol] = 10% (v/v), [PB] = 160 mM, pH = 8.0, $T$ = 37 ℃.

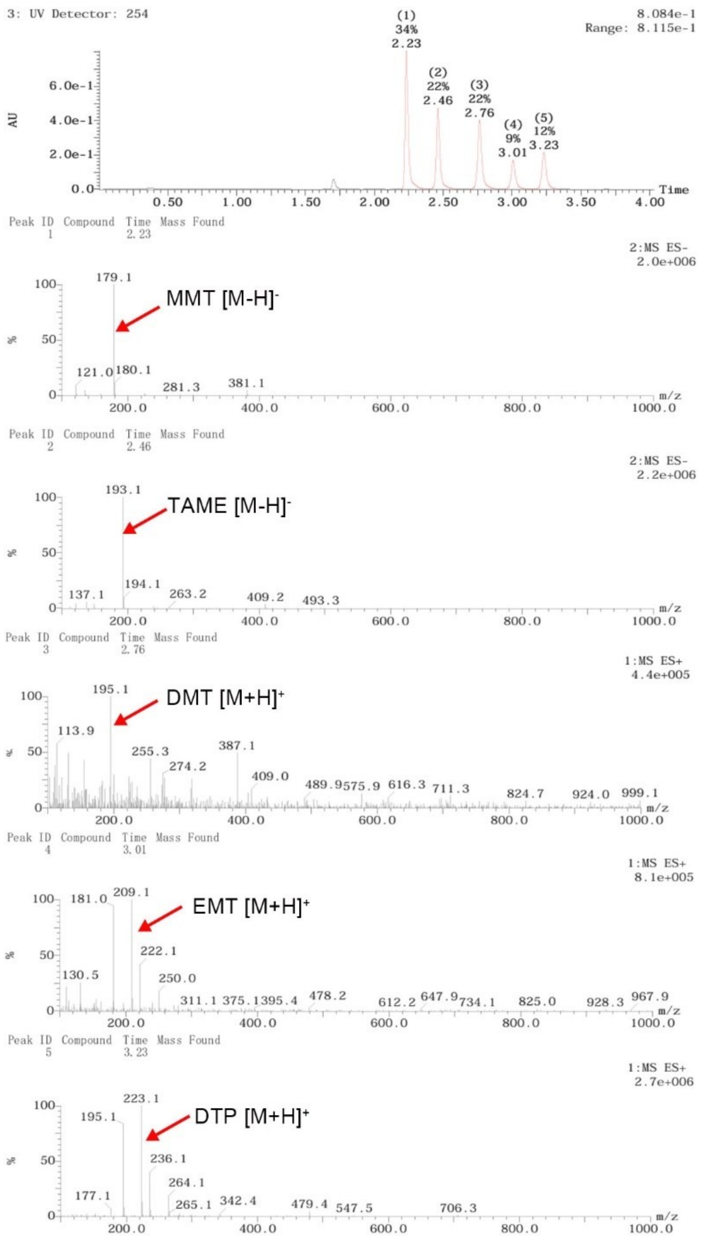


**Fig. S19. LC-MS analysis of the hydrolysis products of DTP catalyzed by Goldenzyme for 24 hours.** Conditions for the hydrolysis: [E] = 0.8 μM, [S] = 5 mM, [methanol] = 10% (v/v), [PB] = 160 mM, pH = 8.0, $T$ = 37 °C.

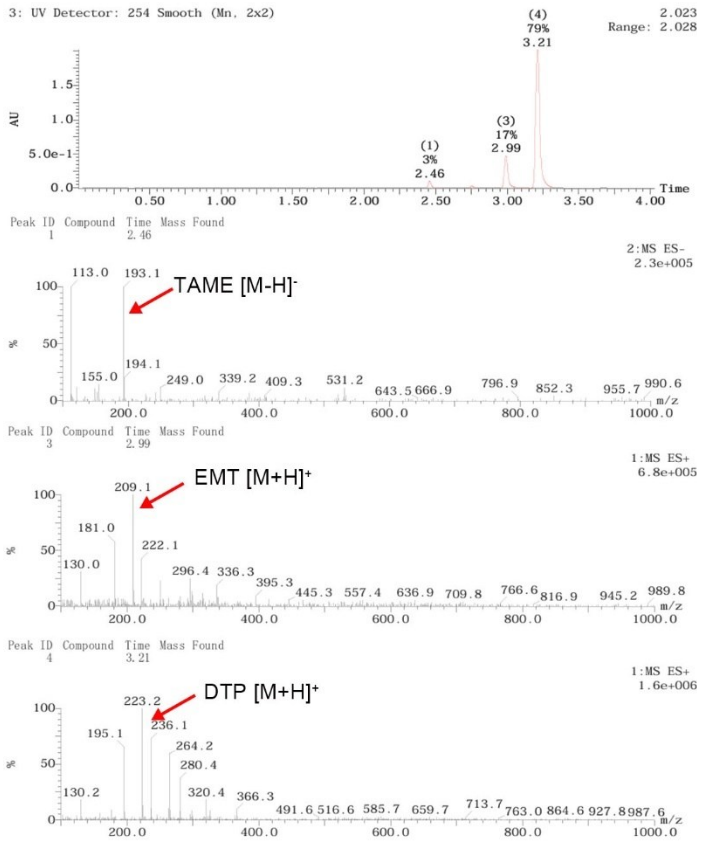


**Fig. S20. LC-MS analysis of the hydrolysis products of DTP catalyzed by the AuNP-Triad5 before CE for 24 hours.** Conditions for the hydrolysis: [E] = 0.8 μM, [S] = 5 mM, [methanol] = 10% (v/v), [PB] = 160 mM, pH = 8.0, $T$ = 37 ℃.

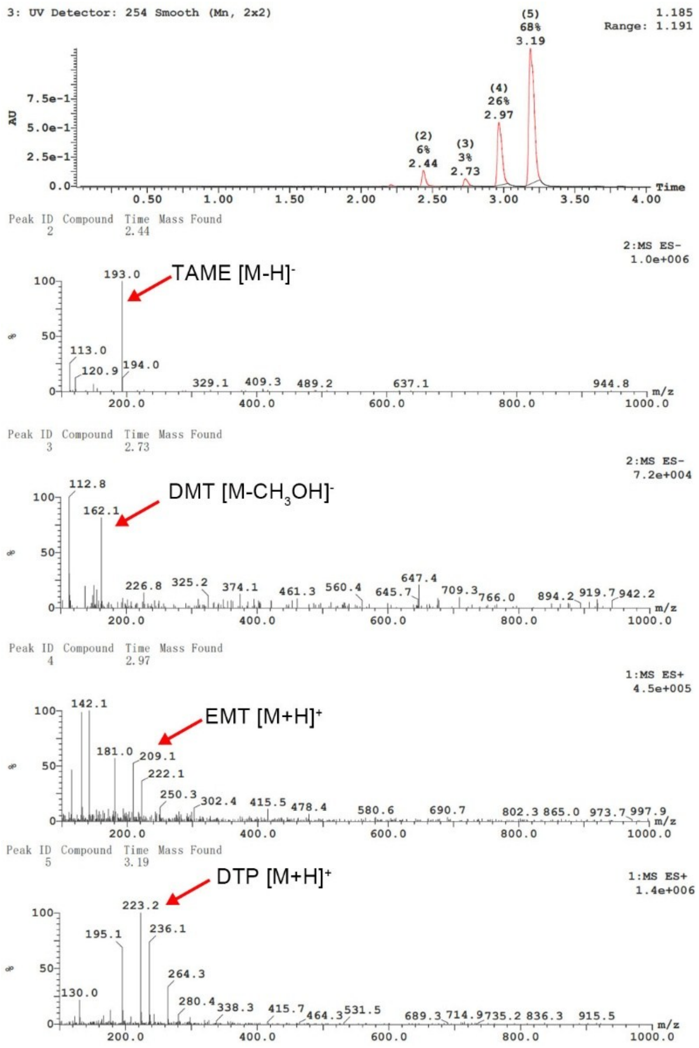


**Fig. S21. LC-MS analysis of the hydrolysis products of DTP catalyzed by the AuNP-Triad5-HSD after CE for 24 hours.** Conditions for the hydrolysis: [E] = 0.8 μM, [S] = 5 mM, [methanol] = 10% (v/v), [PB] = 160 mM, pH = 8.0, $T$ = 37 ℃.

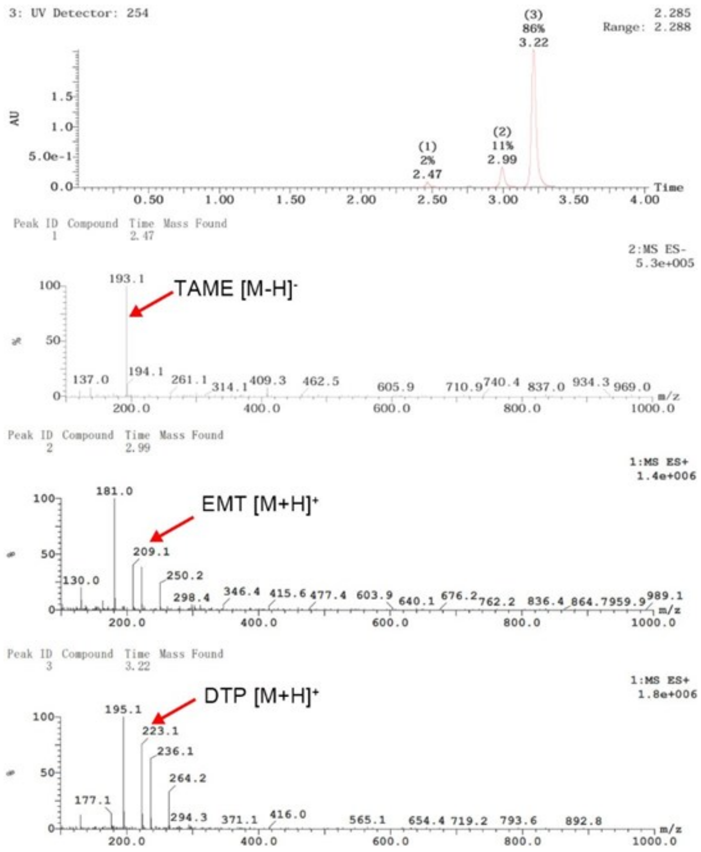


**Fig. S22. LC-MS analysis of the hydrolysis products of DTP catalyzed by the AuNP-CL2 after CE for 24 hours.** Conditions for the hydrolysis: [E] = 0.8 μM, [S] = 5 mM, [methanol] = 10% (v/v), [PB] = 160 mM, pH = 8.0, $T$ = 37 °C.

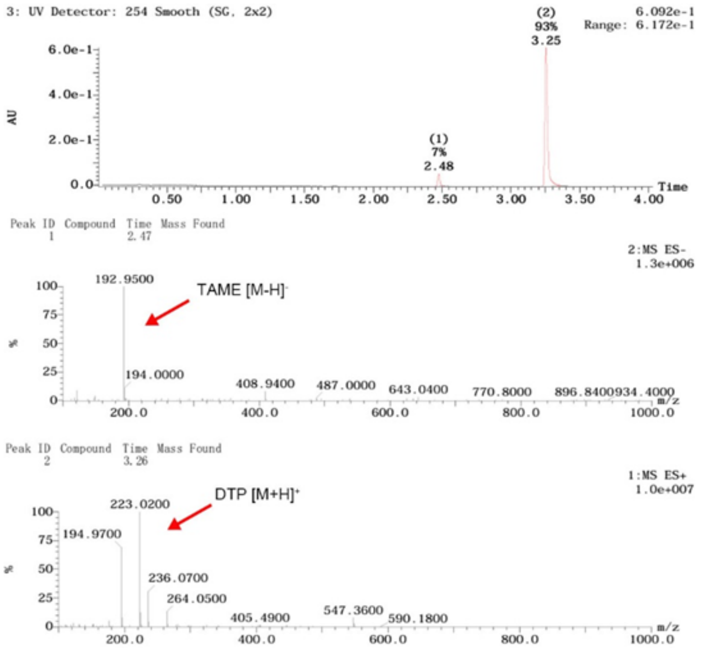


**Fig. S23. LC-MS analysis of the hydrolysis products of DTP in PB buffer after incubation for 48 hours.** Conditions for the hydrolysis: [S] = 5 mM, [acetonitrile] = 10% (v/v), [PB] = 160 mM, pH = 8.0, $T$ = 37 °C.

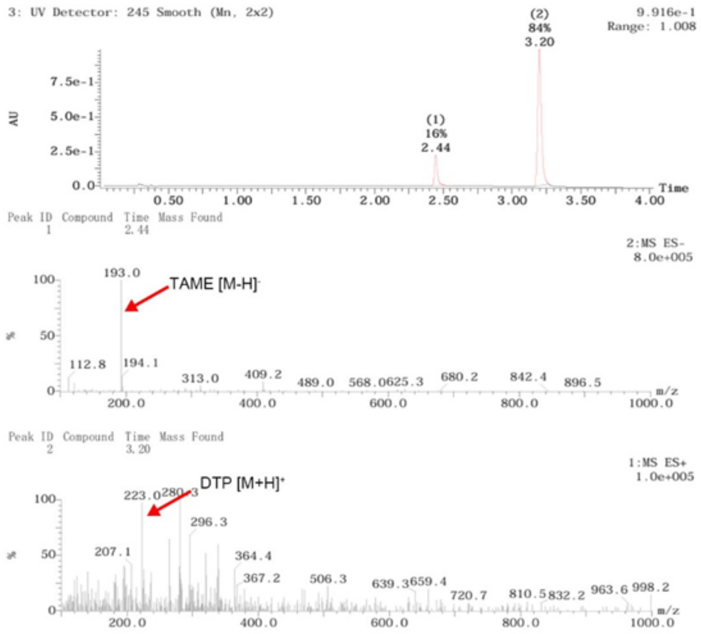


**Fig. S24. LC-MS analysis of the hydrolysis products of DTP in the buffer after CE (the same buffer as that of the Goldenzyme) after incubation for 48 hours.** Conditions for the hydrolysis: [S] = 5 mM, [acetonitrile] = 10% (v/v), [PB] = 160 mM, pH = 8.0, $T$ = 37 °C.

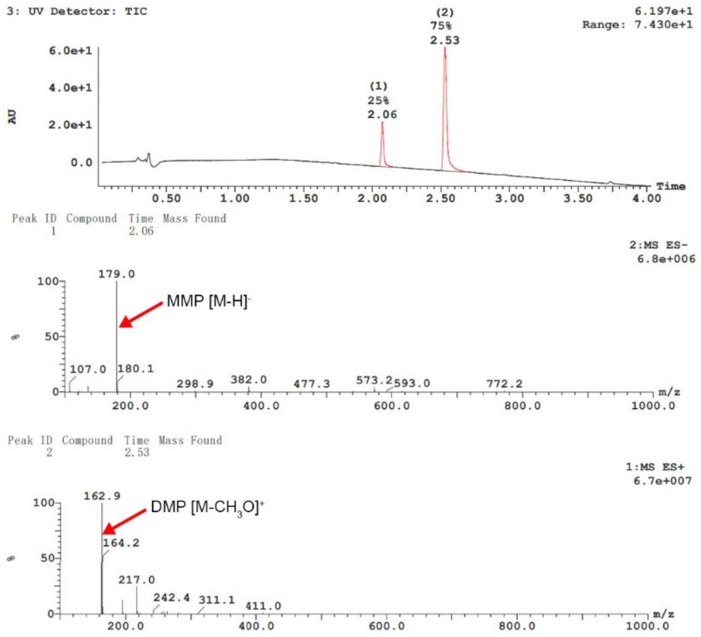


**Fig. S25. LC-MS analysis of the hydrolysis products of DMP catalyzed by α-CT (dissolved in the buffer after CE) for 48 hours.** Conditions for the hydrolysis: [E] = 1.2 μM, [S] = 5 mM, [acetonitrile] = 10% (v/v), [PB] = 160 mM, pH = 8.0, $T$ = 60 °C.

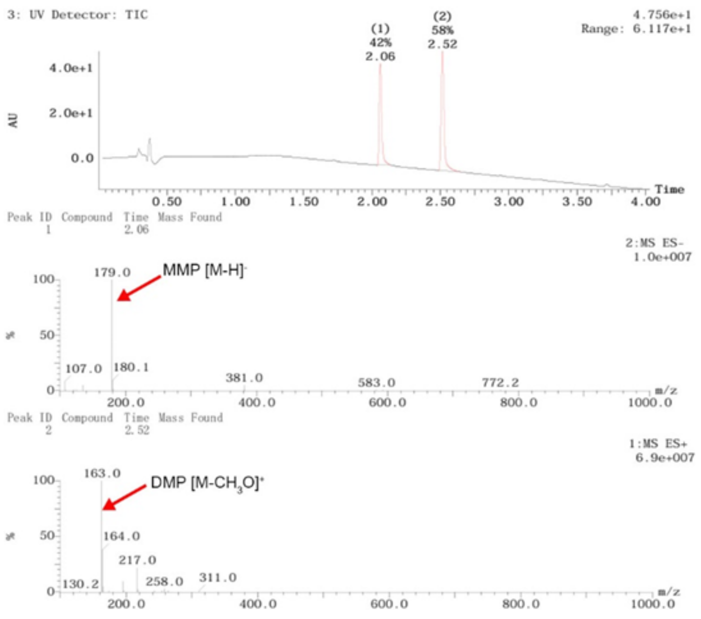


**Fig. S26. LC-MS analysis of the hydrolysis products of DMP catalyzed by Goldenzyme for 48 hours.** Conditions for the hydrolysis: [E] = 1.2 μM, [S] = 5 mM, [acetonitrile] = 10% (v/v), [PB] = 160 mM, pH = 8.0, $T$ = 60 °C.

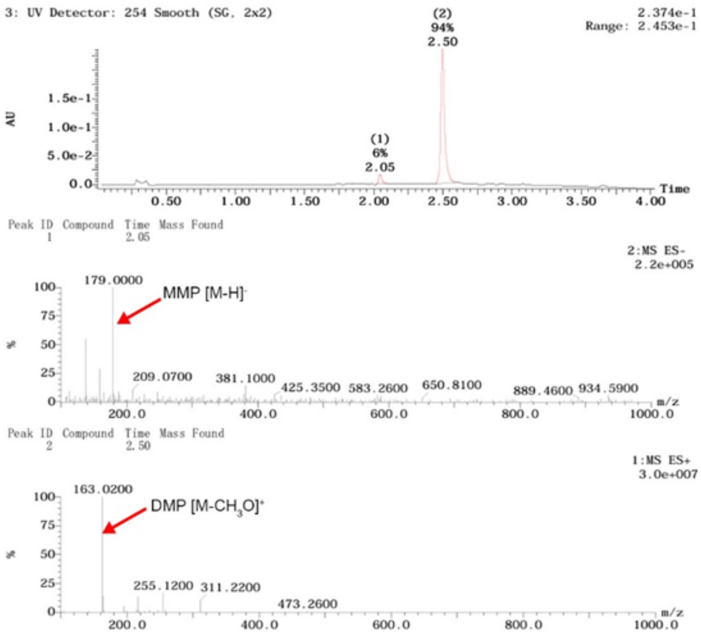


**Fig. S27. LC-MS analysis of the hydrolysis products of DMP catalyzed by the AuNP-Triad5 before CE for 48 hours.** Conditions for the hydrolysis: [E] = 1.2 μM, [S] = 5 mM, [acetonitrile] = 10% (v/v), [PB] = 160 mM, pH = 8.0, $T$ = 60 °C.

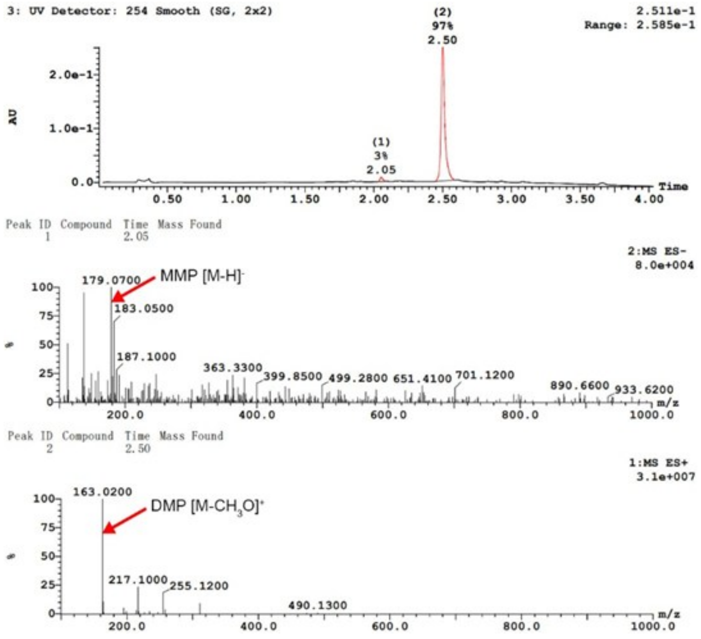


**Fig. S28. LC-MS analysis of the hydrolysis products of DMP catalyzed by the AuNP-Triad5-HSD after CE for 48 hours.** Conditions for the hydrolysis: [E] = 1.2 μM, [S] = 5 mM, [acetonitrile] = 10% (v/v), [PB] = 160 mM, pH = 8.0, $T$ = 60 °C.

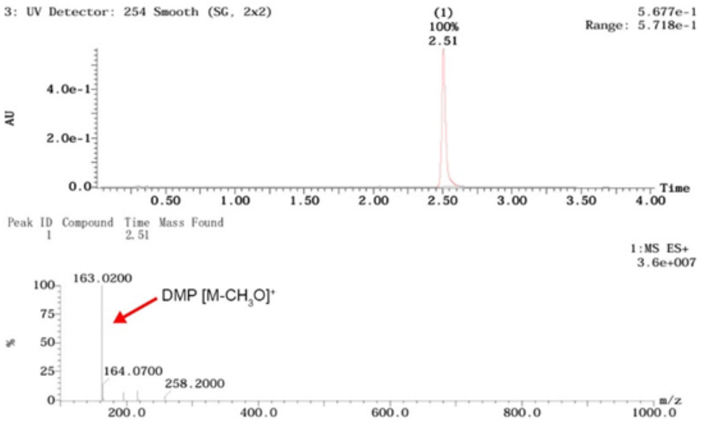


**Fig. S29. LC-MS analysis of the hydrolysis products of DMP catalyzed by the AuNP-CL2 after CE for 48 hours.** Conditions for the hydrolysis: [E] = 1.2 μM, [S] = 5 mM, [acetonitrile] = 10% (v/v), [PB] = 160 mM, pH = 8.0, *T* = 60 °C.

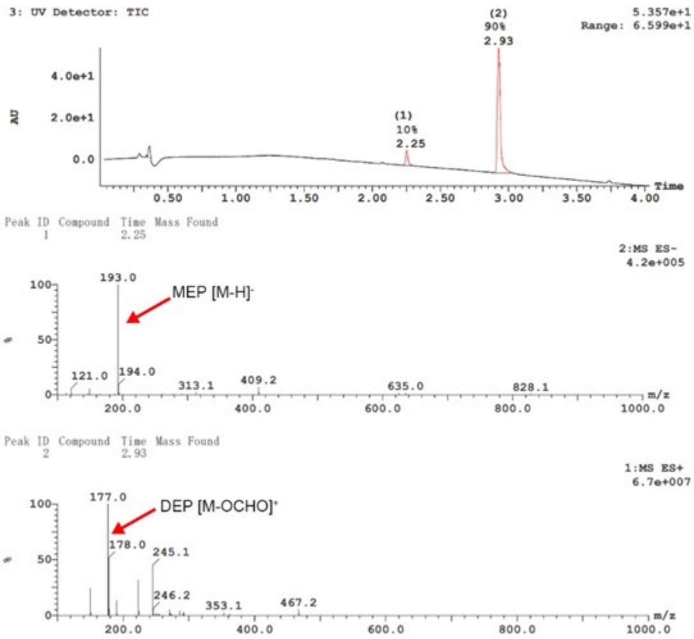


**Fig. S30. LC-MS analysis of the hydrolysis products of DEP catalyzed by α-CT (dissolved in the buffer after CE) for 48 hours.** Conditions for the hydrolysis: [E] = 1.2 μM, [S] = 5 mM, [acetonitrile] = 10% (v/v), [PB] = 160 mM, pH = 8.0, $T$ = 60 °C.

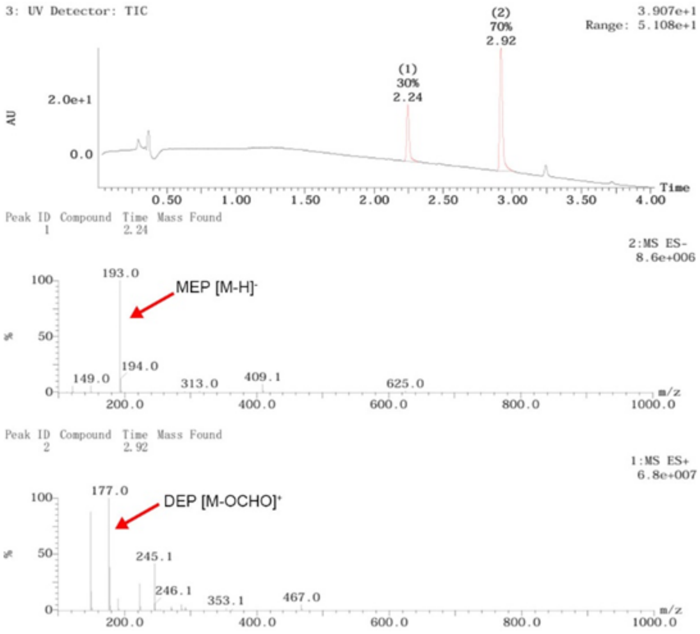


**Fig. S31. LC-MS analysis of the hydrolysis products of DEP catalyzed by Goldenzyme for 48 hours.** Conditions for the hydrolysis: [E] = 1.2 μM, [S] = 5 mM, [acetonitrile] = 10% (v/v), [PB] = 160 mM, pH = 8.0, *T* = 60 °C.

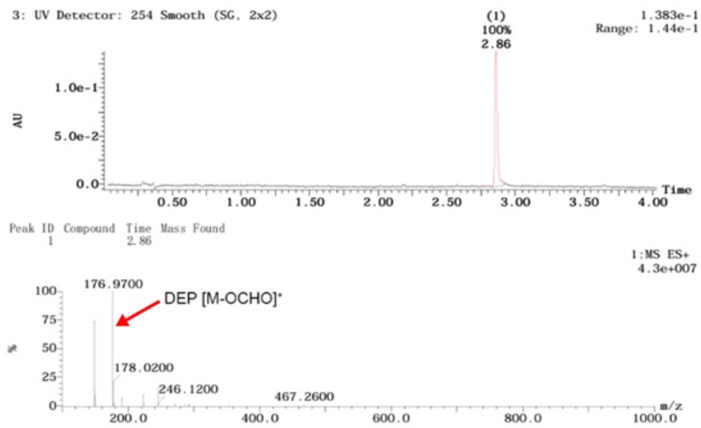


**Fig. S32. LC-MS analysis of the hydrolysis products of DEP catalyzed by the AuNP-Triad5 before CE for 48 hours.** Conditions for the hydrolysis: [E] = 1.2 μM, [S] = 5 mM, [acetonitrile] = 10% (v/v), [PB] = 160 mM, pH = 8.0, $T$ = 60 °C.

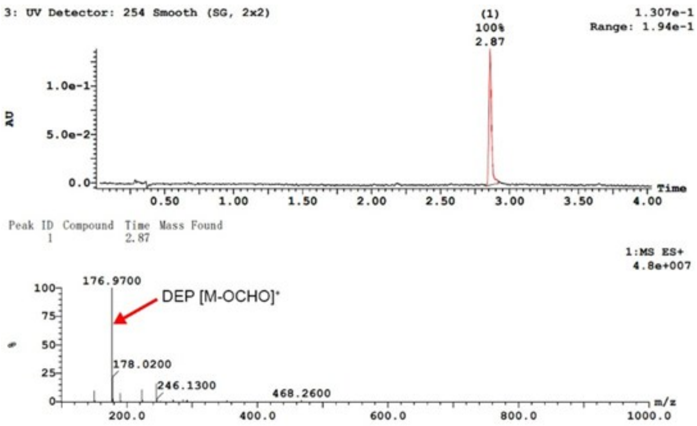


**Fig. S33. LC-MS analysis of the hydrolysis products of DEP catalyzed by the AuNP-Triad5-HSD after CE for 48 hours.** Conditions for the hydrolysis: [E] = 1.2 μM, [S] = 5 mM, [acetonitrile] = 10% (v/v), [PB] = 160 mM, pH = 8.0, $T$ = 60 ℃.

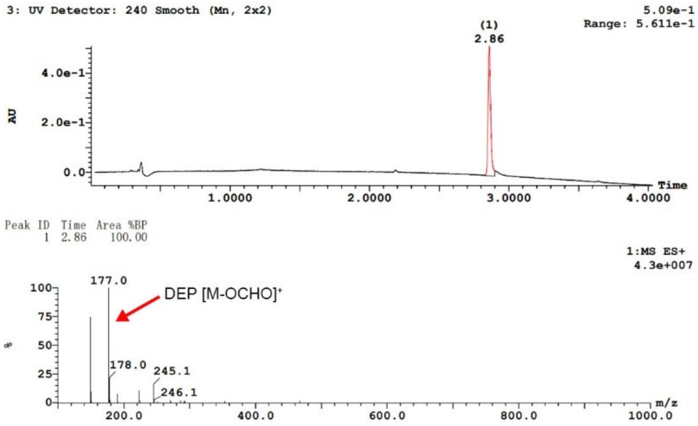


**Fig. S34. LC-MS analysis of the hydrolysis products of DEP catalyzed by the AuNP-CL2 after CE for 48 hours.** Conditions for the hydrolysis: [E] = 1.2 μM, [S] = 5 mM, [acetonitrile] = 10% (v/v), [PB] = 160 mM, pH = 8.0, $T$ = 60 ℃.

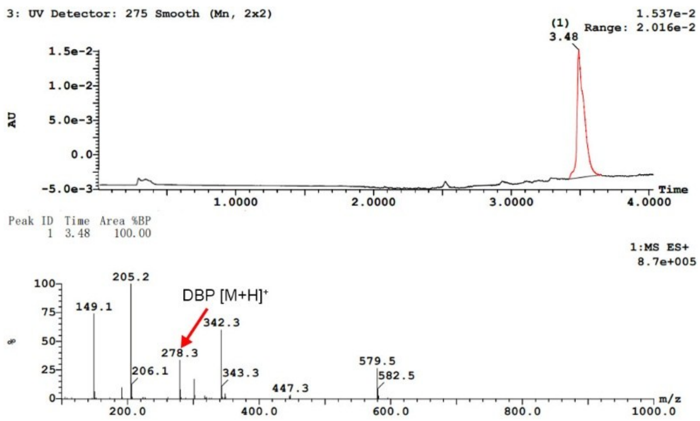


**Fig. S35. LC-MS analysis of the hydrolysis products of DBP catalyzed by α-CT (dissolved in the buffer after CE) for 48 hours.** Conditions for the hydrolysis: [E] = 1.2 μM, [S] = 5 mM, [acetonitrile] = 10% (v/v), [PB] = 160 mM, pH = 8.0, $T$ = 60 °C.

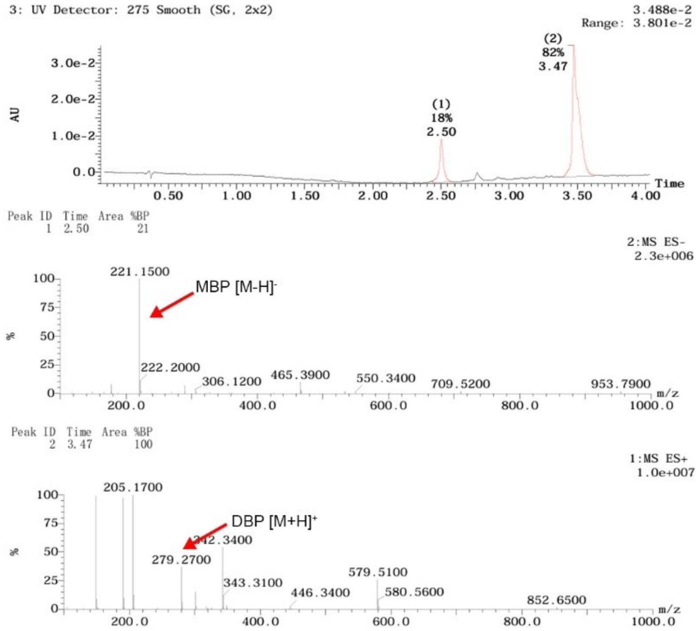


**Fig. S36. LC-MS analysis of the hydrolysis products of DBP catalyzed by Goldenzyme for 48 hours.** Conditions for the hydrolysis: [E] = 1.2 μM, [S] = 5 mM, [acetonitrile] = 10% (v/v), [PB] = 160 mM, pH = 8.0, $T$ = 60 °C.

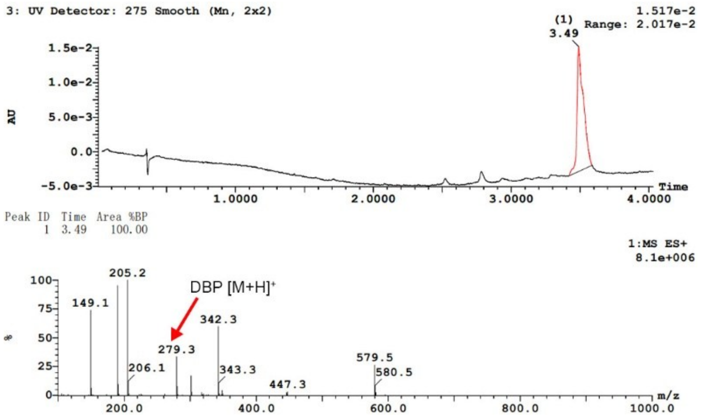


**Fig. S37. LC-MS analysis of the hydrolysis products of DBP catalyzed by the AuNP-Triad5 before CE for 48 hours.** Conditions for the hydrolysis: [E] = 1.2 μM, [S] = 5 mM, [acetonitrile] = 10% (v/v), [PB] = 160 mM, pH = 8.0, $T$ = 60 ℃.

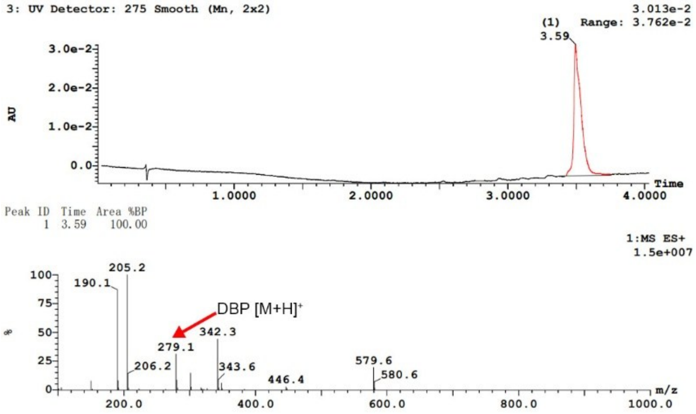


**Fig. S38. LC-MS analysis of the hydrolysis products of DBP catalyzed by the AuNP-Triad5-HSD after CE for 48 hours.** Conditions for the hydrolysis: [E] = 1.2 μM, [S] = 5 mM, [acetonitrile] = 10% (v/v), [PB] = 160 mM, pH = 8.0, $T$ = 60 ℃.

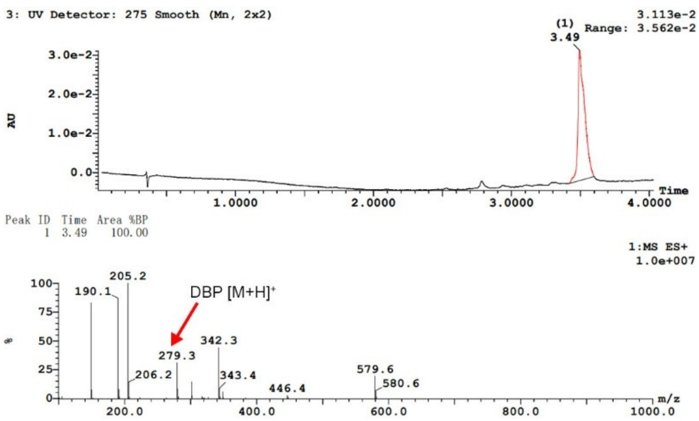


**Fig. S39. LC-MS analysis of the hydrolysis products of DBP catalyzed by the AuNP-CL2 after CE for 48 hours.** Conditions for the hydrolysis: [E] = 1.2 μM, [S] = 5 mM, [acetonitrile] = 10% (v/v), [PB] = 160 mM, pH = 8.0, $T$ = 60 °C.

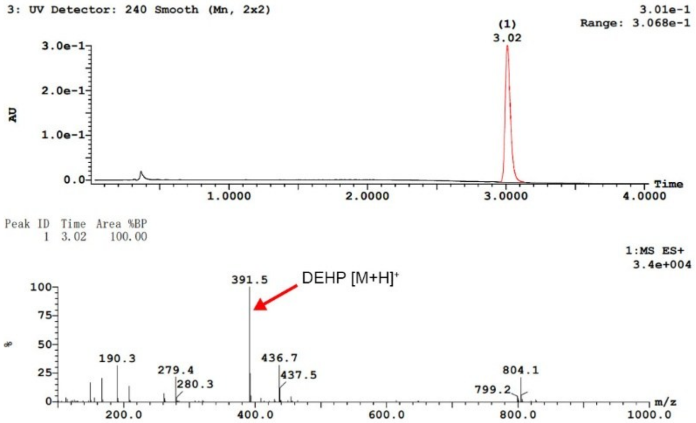


**Fig. S40. LC-MS analysis of the hydrolysis products of DEHP catalyzed by α-CT (dissolved in the buffer after CE) for 72 hours.** Conditions for the hydrolysis: [E] = 1.2 μM, [S] = 5 mM, [acetonitrile] = 10% (v/v), [PB] = 160 mM, pH = 8.0, $T$ = 60 °C.

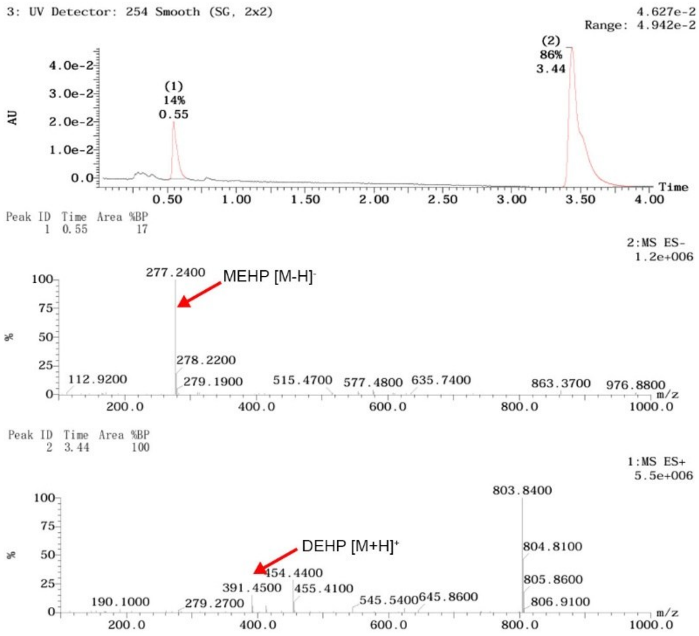


**Fig. S41. LC-MS analysis of the hydrolysis products of DEHP catalyzed by Goldenzyme for 72 hours.** Conditions for the hydrolysis: [E] = 1.2 μM, [S] = 5 mM, [acetonitrile ] = 10% (v/v), [PB] = 160 mM, pH = 8.0, $T$ = 60 °C.

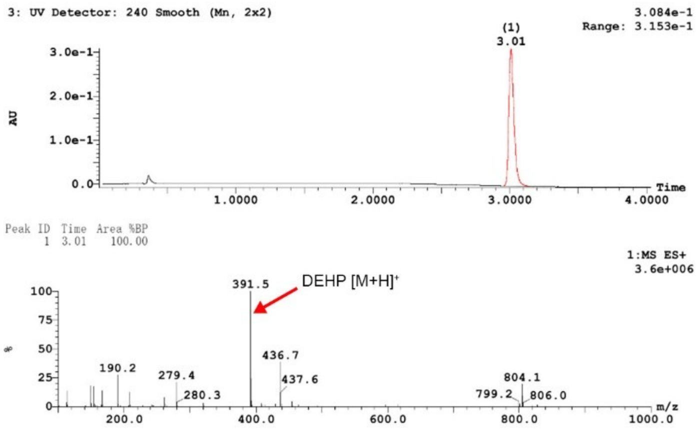


**Fig. S42. LC-MS analysis of the hydrolysis products of DEHP catalyzed by the AuNP-Triad5 before CE for 72 hours.** Conditions for the hydrolysis: [E] = 1.2 μM, [S] = 5 mM, [acetonitrile ] = 10% (v/v), [PB] = 160 mM, pH = 8.0, $T$ = 60 °C.

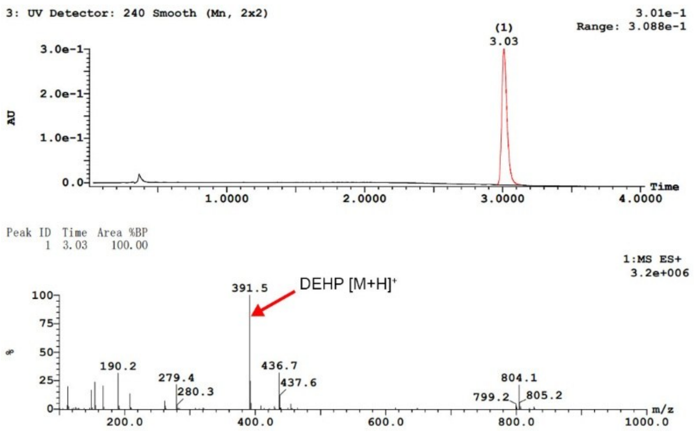


**Fig. S43. LC-MS analysis of the hydrolysis products of DEHP catalyzed by the AuNP-Triad5-HSD after CE for 72 hours.** Conditions for the hydrolysis: [E] = 1.2 μM, [S] = 5 mM, [acetonitrile ] = 10% (v/v), [PB] = 160 mM, pH = 8.0, $T$ = 60 °C.

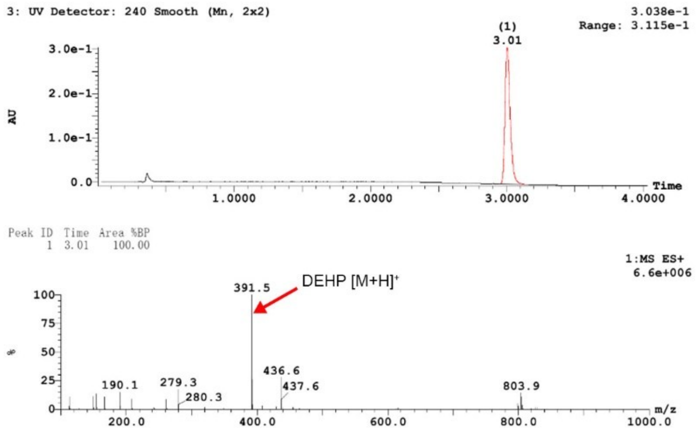


**Fig. S44. LC-MS analysis of the hydrolysis products of DEHP catalyzed by the AuNP-CL2 after CE for 72 hours.** Conditions for the hydrolysis: [E] = 1.2 μM, [S] = 5 mM, [acetonitrile] = 10% (v/v), [PB] = 160 mM, pH = 8.0, $T$ = 60 °C.

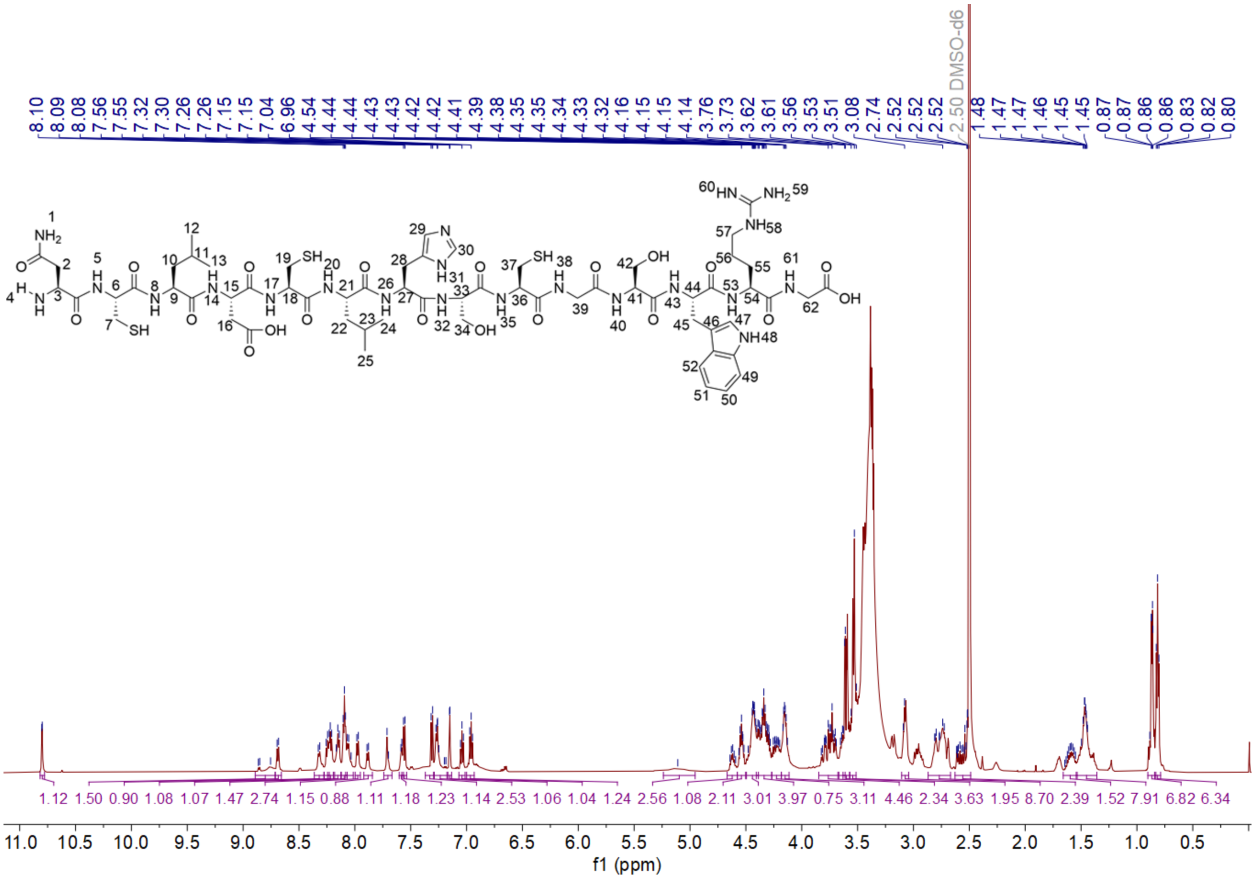

**Fig. S45. $^{1}$H NMR spectra of Triad5 recorded at 25 °C in DMSO-d6.** Assignment of the hydrogen peaks of Triad5 (600 MHz, δ in ppm): 10.80 (d, $J$ = 2.4 Hz, 1H, H-48), 8.75 (s, 1H, H-31), 8.69 (d, $J$ = 7.7 Hz, 1H, H-14), 8.32 (d, $J$ = 7.9 Hz, 1H, H-26), 8.25 (d, $J$ = 7.7 Hz, 2H, H-35, H-43), 8.24 – 8.19 (m, 1H, H-38), 8.18 – 8.12 (m, $J$ = 7.7 Hz, 2H, H-5, 53), 8.11 – 8.10 (m, 1H, H-17), 8.10 – 8.08 (m, 1H, H-32), 8.08 – 8.07 (m, 1H, H-61), 8.06 (d, $J$ = 7.5 Hz, 1H, H-8), 7.98 (d, $J$ = 7.8 Hz, 1H, H-20), 7.88 (d, $J$ = 8.0 Hz, 1H, H-40), 7.71 (s, 1H, H-30), 7.58 (s, 1H, H-29), 7.56 (d, $J$ = 8.0 Hz, 1H, H-52), 7.31 (d, $J$ = 8.1 Hz, 1H, H-49), 7.27 (2H, s, H-1), 7.26 (s, 1H, H-60), 7.15 (d, $J$ = 2.4 Hz, 1H, H-47), 7.04 (t, $J$ = 7.5 Hz, 1H, H-50), 6.96 (t, $J$ = 7.4 Hz, 1H, H-51), 5.11 (s, 2H, H-59), 4.68 – 4.62 (m, 1H, H-44), 4.58 – 4.50 (m, 2H, H-21, 54), 4.49 – 4.39 (m, 3H, H-15, 36, 18), 4.41 – 4.27 (m, 4H, H-27, 41, 57), 4.23 (ddd, $J$ = 18.2, 9.6, 6.4 Hz, 1H, H-9), 4.15 (m, 3H, H-6, 39), 3.85 – 3.68 (m, 5H, H-16, 33, 62), 3.63 – 3.57 (m, 1H, H-3), 3.61 (d, $J$ = 3.5 Hz, 2H, H-4), 3.53 (m, 4H, H-34, 42), 3.09 (d, $J$ = 6.8 Hz, 2H, H-45), 2.87 – 2.67 (m, 8H, H-7, 19, 28, 37), 2.63 – 2.49 (m, 3H, H-2, 58), 1.60 (m, 2H, H-11, 23), 1.54 – 1.36 (m, 8H, H-10, 22, 55, 56), 0.87 (d, $J$ = 6.6, 6H, H-24, 25), 0.82 (d, $J$ = 7.1 Hz, 6H, H-12, 13).

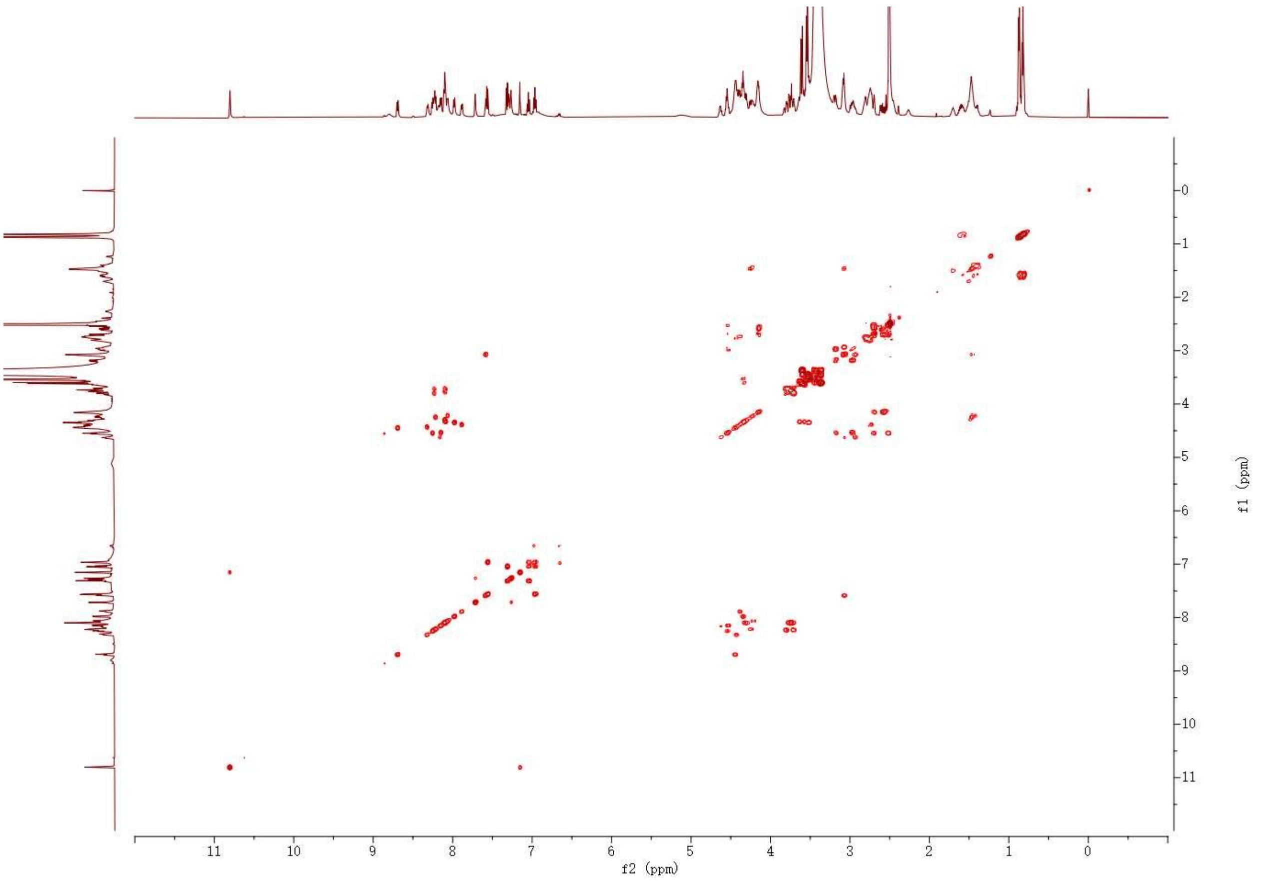


**Fig. S46. 1H-1H COSY spectrum of free peptide Triad5 before deuterium exchange recorded at 25 °C in DMSO-d6.**

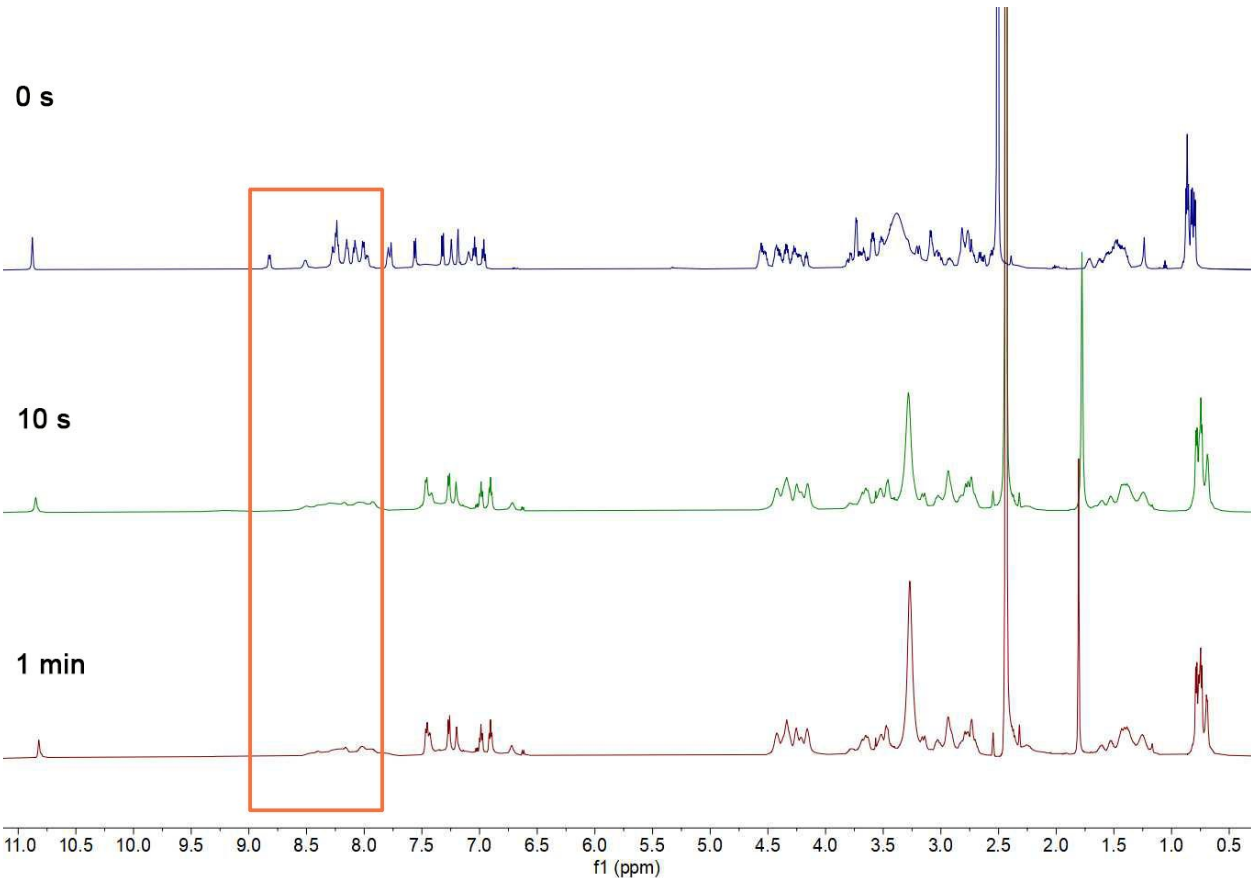


**Fig. S47. $^{1}H$ NMR spectra of the free peptide Triad5 after hydrogen-deuterium exchange in $D_2O$ for certain periods of time.** The spectra were recorded at 25 °C using lyophilized powder re-dissolved in DMSO-d6. The orange red rectangle shows the peaks of most backbone amide hydrogens.

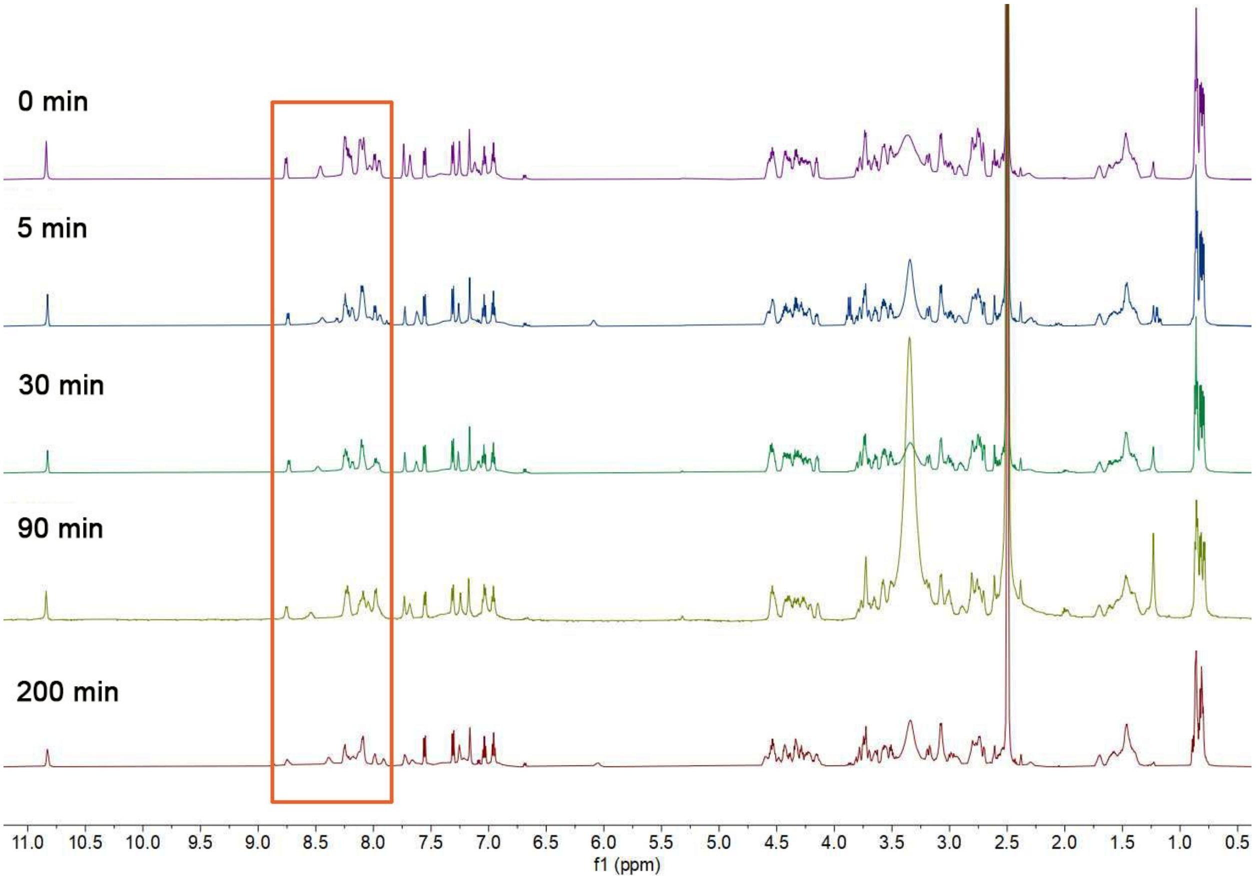


**Fig. S48. $^1$H NMR spectra of the Triad5 in 50% TFE after hydrogen-deuterium exchange in 50% $D_2O$ plus TFE-d3 for certain periods of time.** The spectra were recorded at 25 °C using lyophilized powder re-dissolved in DMSO-d6. The orange red rectangle shows the peaks of most backbone amide hydrogens.

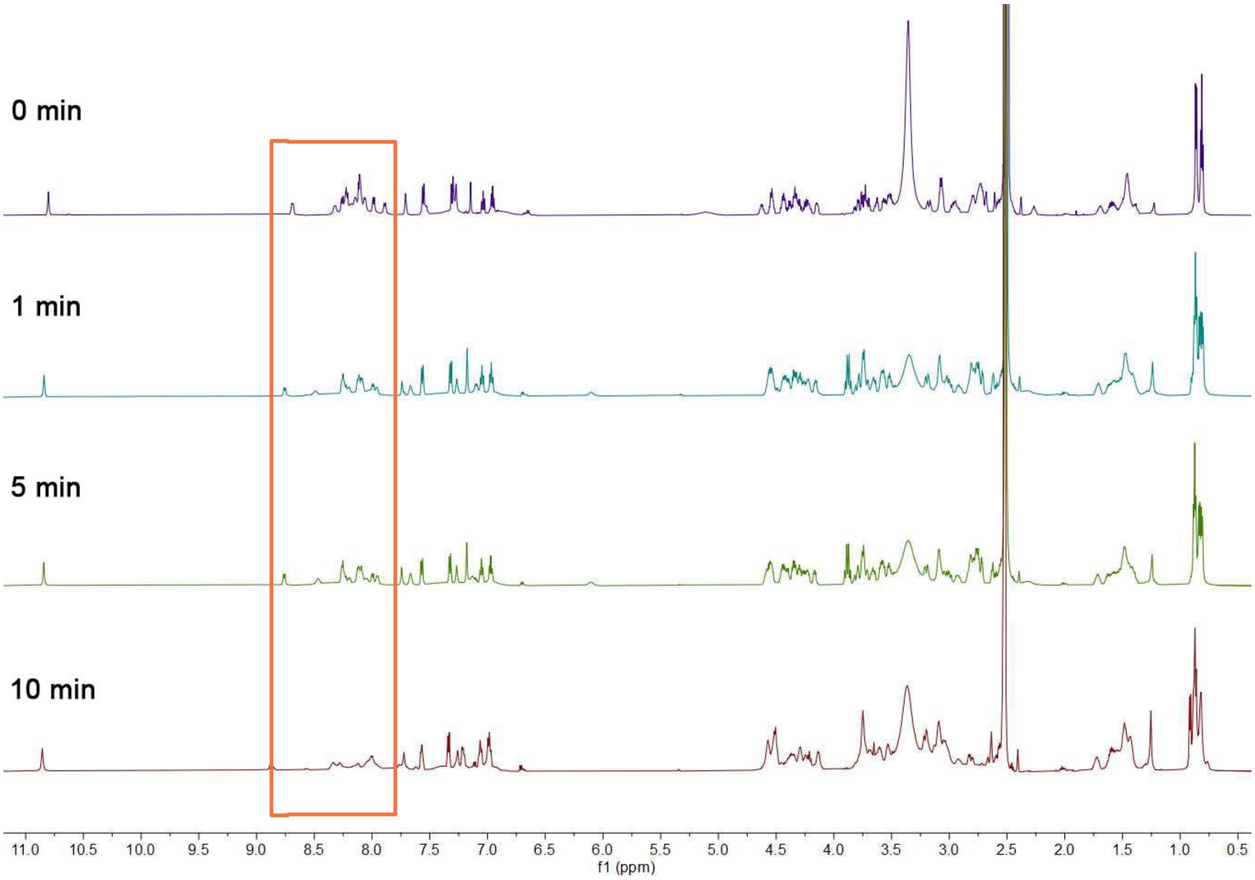


**Fig. S49. [1]H NMR spectra of the AuNP-Triad5 before CE after hydrogen-deuterium exchange in $D_2O$ for certain periods of time.** The spectra were recorded at 25 °C using lyophilized powder re-dissolved in DMSO-d6. The orange red rectangle shows the peaks of most backbone amide hydrogens.

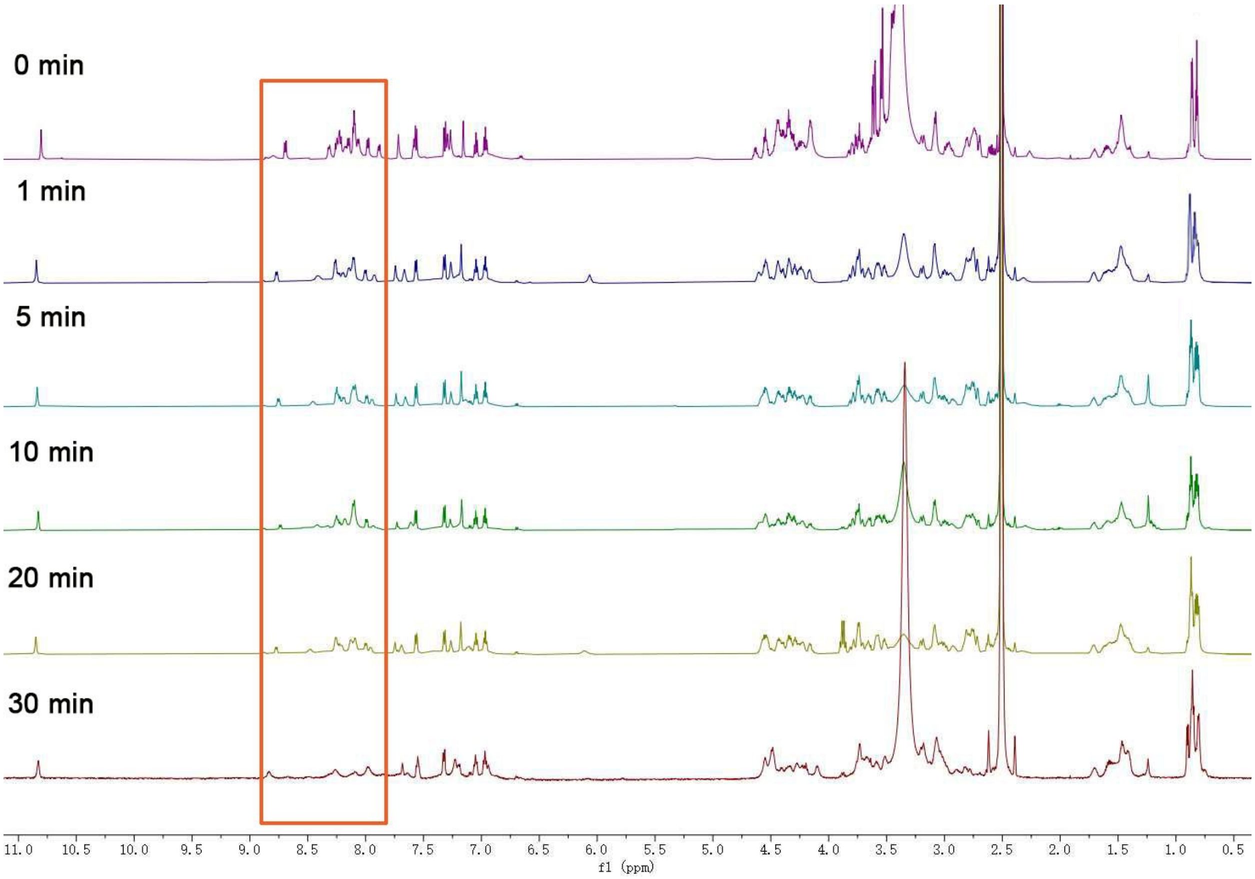


**Fig. S50. [1]H NMR spectra of the AuNP-Triad5 after CE (Goldenzyme) after hydrogen-deuterium exchange in $D_2O$ for certain periods of time.** The spectra were recorded at 25 °C using lyophilized powder re-dissolved in DMSO-d6. The orange red rectangle shows the peaks of most backbone amide hydrogens.

**Table S1. Sequences and properties of the designed peptides**

| **Peptide** | **Peptide sequence** | ***p*I** | **MW (Da)** | **$\varepsilon_{280\ nm}$ ($M^{-1}\cdot cm^{-1}$)** |
|---|---|---|---|---|
| Triad5 | NCLDCLHSCGSWRG | 6.72 | 1550.74 | 5625 |
| Triad5-D | NCL**A**CLHSCGSWRG | 7.98 | 1506.74 | 5625 |
| Triad5-H | NCLDCL**A**SCGSWRG | 5.82 | 1484.68 | 5625 |
| Triad5-S | NCLDCLH**A**CG**A**WRG | 6.72 | 1518.75 | 5625 |
| Triad5-HSD | NCL**A**CL**AA**CG**A**WRG | 7.97 | 1408.67 | 5625 |
| Triad5-O | NCLDCLHSCGSW | 5.08 | 1337.51 | 5625 |
| Triad5~W | **W**CLDCLHSCGS**N**RG | 6.72 | 1550.74 | 5625 |
| CL2 | GYCGTNPNYFSCD | 3.80 | 1440.52 | 3104 |

**Table S2. Effect of peptide density on the initial catalytic rate of Goldenzyme (after subtracting the rate of AuNP-CL2 under the corresponding conditions).** Conditions for the hydrolysis of *p*-NPA: [AuNP] = 0.5 μM, [*p*-NPA] = 1 mM, [acetonitrile ] = 2% (v/v), [PB] = 40 mM, pH = 7.0, $T$ = 25 °C. ($v_{total}$: catalytic rate per particle; $v_{single}$: catalytic rate per peptide)

| **peptides per AuNP** | $v_{total}$ **(μM·s$^{-1}$)** | $v_{single}$ **(μM·s$^{-1}$)** |
|---|---|---|
| 10 | 5.1 ± 0.5 | 0.51 ± 0.05 |
| 20 | 18.0 ± 0.3 | 0.90 ± 0.02 |
| 30 | 23.5 ± 2.4 | 0.78 ± 0.08 |
| 40 | 22.1 ± 6.2 | 0.55 ± 0.15 |

**Table S3. Initial catalytic rates ($v_0$) of the AuNP-peptide conjugates before and after CE and the corresponding free peptides for the hydrolysis of *p*-NPA (after subtracting the rate of AuNP-CL2 under the corresponding conditions).** Conditions for the hydrolysis: [AuNP-peptide] = 0.5 μM, [Peptide] = 10 μM, [S] = 1 mM, [acetonitrile] = 2% (v/v), [PB] = 40 mM, pH = 7.0, $T$ = 25 °C.

| **Peptide** | **$v_0$ (μM·s$^{-1}$)** | | |
|---|---|---|---|
| | **AuNP-peptide after CE** | **AuNP-peptide before CE** | **Free peptide** |
| **Triad5** | 18.2 ± 3.5 | 6.5 ± 1.3 | 0.013 ± 0.004 |
| **Triad5-D** | 1.0 ± 0.1 | 2.1 ± 1.3 | 0.016 ± 0.001 |
| **Triad5-H** | 3.1 ± 1.9 | 2.2 ± 0.2 | 0.015 ± 0.002 |
| **Triad5-S** | 8.6 ± 1.4 | 3.2 ± 0.3 | 0.012 ± 0.003 |
| **Triad5-HSD** | 0.1 ± 0.1 | 0.2 ± 0.3 | 0.014 ± 0.003 |
| **Triad5-O** | 0.1 ± 0.1 | 0.9 ± 0.2 | 0.010 ± 0.008 |
| **Triad5~W** | 0.1± 0.7 | 0.1 ± 0.4 | 0.016 ± 0.004 |
